# Inflammatory burning pain depends on a specific line of nociceptors

**DOI:** 10.64898/2026.09.14.748479

**Authors:** Muhammad Rizki Febrianto, Deemah Aldossary, David Zimmermann, Jahanzaib Irfan, Michelle Y Meng, Ilaria Sanvido, Luming Zhou, Guiping Kong, Angela Kecskes, Paolo La Montanara, Jose Vicente Torres-Perez, Krisztina Deak-Pocsai, Botond Gaal, Angelina Mira D’Ercole, Nicole Li, Eva Szoke, Jianglin Wang, Klara Matesz, Simon Arthur, Erika Pinter, Simone di Giovanni, Michaela Kress, Istvan Nagy

**Author notes:** contributed equally (Muhammad Rizki Febrianto, Deemah Aldossary, David Zimmermann). contributed equally (Angelina Mira D’Ercole, Nicole Li). contributed equally (Michaela Kress, Istvan Nagy). Correspondence: Istvan Nagy, Department of Surgery and Cancer Imperial College London Hammersmith Hospital Campus Du Cane Road, London W12 0NN United Kingdom.

## Abstract

The development and persistence of burning pain and heat hyperalgesia following tissue injury and the subsequent inflammatory response, depend on the nuclear enzyme mitogen-and stress-activated kinase 1 (MSK1) expressed in a specific subset of transient receptor potential subfamily V member 1 (TRPV1)-expressing primary sensory neurons termed nociceptors, which are specialised for detecting harmful stimuli. Inflammation up-regulates and activates MSK1, and MSK1 governs TRPV1 expression in the MSK1 and TRPV1 co-expressing mouse and human nociceptors. Importantly, inhibition of inflammatory MSK1-mediated TRPV1 upregulation protects from heat hypersensitivity without affecting other relevant TRPV1 functions, such as the sensation of acute painful heat stimuli or the maintenance of the body core temperature. The newly discovered importance of the interaction between MSK1 and TRPV1 in a specific line of nociceptors provides mechanistic understanding of inflammatory pain and heat hyperalgesia pathogenesis.

---

Inflammation is generally associated with enduring tenderness and burning pain (heat hyperalgesia) of the affected tissue, and this inflammatory pain persists after tissue injury (Basbaum et al., 2009; Parisien et al., 2022). A major group of primary afferent sensory neurons specifically detect noxious stimuli, and these nociceptors are pivotal for the development and persistence of inflammatory pain (Berta et al., 2023; Patapoutian et al., 2009). Almost twenty transcriptionally distinct sub-populations of primary sensory neurons exhibiting remarkable correlation between transcriptional, morphological and functional properties are emerging from transcriptomic profiling (Bhuiyan et al., 2024; Nguyen et al., 2021; Qi et al., 2024; Renthal et al., 2020; Wang et al., 2021). The existence of differing sensory neuron sub-populations indicates cellular specificity for various somatosensory modalities including responsiveness to specific noxious stimuli that characterises different subgroups of nociceptors.

About 50 % of nociceptors express the multimodal heat transducer ion channel transient receptor potential subfamily V member 1 (TRPV1) which is critically involved in inflammatory heat hyperalgesia (Baiou et al., 2007; Caterina et al., 2000; Caterina et al., 1997; Davis et al., 2000; Guo et al., 1999; Nagy et al., 2014). TRPV1 becomes hypersensitive to heat stimuli following inflammatory mediators-induced activation of specific metabotropic or ionotropic receptors at the peripheral nociceptor terminal and their downstream signaling cascades involving protein kinases and phosphorylation of TRPV1 at serine, threonine or tyrosine residues (Nagy et al., 2014; Zimmermann et al., 2025). The level of phosphorylation determines the channel’s function, sensitivity to thermal stimuli, open probability, and membrane insertion (Andratsch et al., 2009; Bhave et al., 2002; Bonnington & McNaughton, 2003; Premkumar & Ahern, 2000; Prescott & Julius, 2003; Rathee et al., 2002; Zhang et al., 2005; Zhang et al., 2008). In addition, kinases including the stress-activated protein kinase 2 (SAPK2)/p38 within the mitogen-activated protein kinase (MAPK) pathway, upregulate *TRPV1* expression and the number of available channel proteins, which is critically important for the persistence of inflammatory heat hyperalgesia (Ji & Woolf, 2001). TRPV1, in addition to inflammatory heat hyperalgesia, also contributes to vital functions, such as the protective withdrawal reflex from acute noxious heat stimuli of healthy tissues and maintenance of the body core temperature (Vandewauw et al., 2018; Yue et al., 2022). These important roles accounting for severe unwanted effects such as fever and increased risk of attaining burn injury have disqualified the clinical use of TRPV1 inhibitors for inflammatory heat hyperalgesia (Chizh et al., 2007; Gavva, 2008; Gavva et al., 2008; Szallasi & Sheta, 2012).

The nuclear mitogen-and stress-activated protein kinases (MSK) 1 and MSK2 contribute to various adaptive responses including inflammatory hyperalgesia *but not* acute heat pain (Blum et al., 1999; Brami-Cherrier et al., 2005; Chwang et al., 2007; Privitera et al., 2020; Sindreu et al., 2007; Torres-Perez et al., 2017; Zhou et al., 2019). MSKs are activated by MAPKs including p38 and activate transcriptional regulators such as cAMP response element binding protein (CREB), nuclear factor kappa-light-chain-enhancer of activated B cells (NF-κB) and histone 3 (H3) at serine (S) 10 and S28 (Chen et al., 2005; Deak et al., 1998; Soloaga et al., 2003; Wiggin et al., 2002). Based on these functions we hypothesized that MSKs may regulate *TRPV1* expression in nociceptor types that are essential for inflammatory heat hyperalgesia. Therefore, we explored which of the MSKs defined functionally distinct sub-populations or states of nociceptors, responding to inflammation with upregulation of TRPV1 for the induction and persistence of inflammatory heat hyperalgesia.

## Results

### MSK1^-/-^mice protected from inflammatory heat hypersensitivity

To identify which MSK isoform(s) were predominantly involved in the development of inflammatory heat hypersensitivity, we first assessed pain-related behaviour of MSK1^-/-^and MSK2^-/-^mice in two inflammatory pain models (Figure 1A-D). In the Hargreaves test, which assesses sensitivity to heat, naive MSK1^-/-^and MSK2^-/-^mice performed similarly to naive wild type (WT) indicating that depletion of neither MSK1 nor MSK2 compromises acute noxious heat detection in healthy tissues (Figure 1A, B). After carrageenan injection (Abboud et al., 2021), WT mice became hypersensitive to heat stimulation, and the hypersensitivity did not recover until the end of the observation period. In contrast, MSK1^-/-^mice but not MSK2^-/-^mice were largely protected from heat hypersensitivity (Figure 1A). No changes were observed on the contralateral side or upon vehicle injection in mice of any genotype (Supplementary Figure 1A, B). Induction of inflammation by Complete Freund’s Adjuvant (CFA) (Abboud et al., 2021) resulted in maintained hypersensitivity in both WT and MSK2^-/-^mice, whereas MSK1^-/-^mice were fully protected at all time points (Figure 1B).

**Figure 1:**
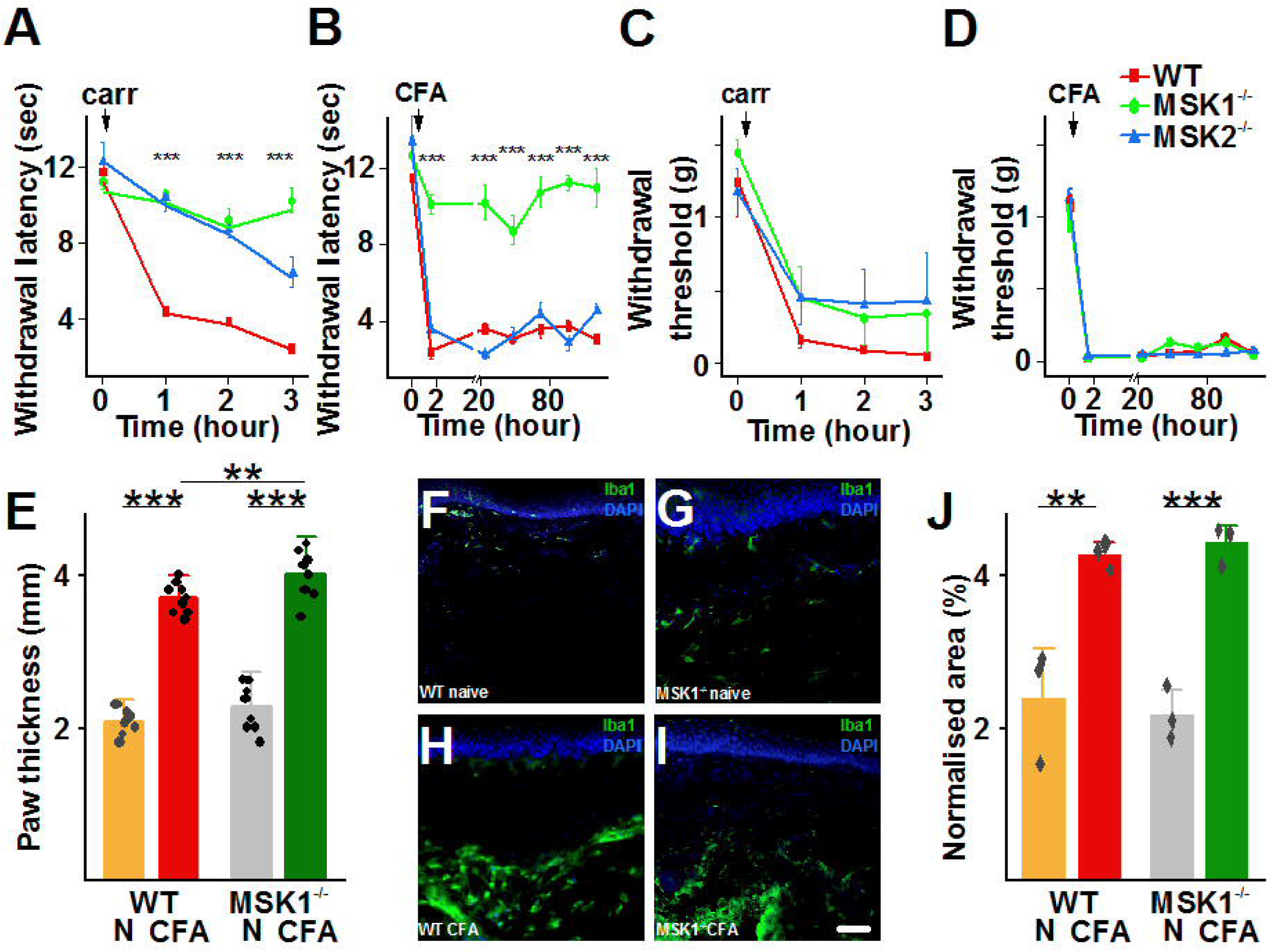
Global MSK1 deletion inhibits the development of inflammatory heat hypersensitivity without affecting noxious heat detection in healthy tissues or attenuating inflammation (A) and (B) Paw withdrawal latency of the ipsilateral hind paw in WT, MSK1^-/-^and MSK2^-/-^ mice, after unilateral injection the inflammation-inducing agents, carrageenan (carr; (A)) or Complete Freund’s Adjuvant (CFA; (B)) into the left paw. Global deletion of MSK1, but not MSK2 prevented inflammation-induced reduction in paw withdrawal latency, which indicates that MSK1^-/-^ mice were protected from the development inflammatory heat hypersensitivity (two-way repeated measures ANOVA, Bonferroni post-hoc test, *** indicate p<0.0001 between values of WT and MSK1-/- mice; n= 5-7). (C) and (D) Paw withdrawal threshold of the ipsilateral hind paw in WT, MSK1^-/-^ and MSK2^-/-^ mice after injecting carrageenan (carr; (C)) or CFA (D) into the left hind paw. Deletion of neither MSK1 or MSK2 had any effect on the paw withdrawal threshold either before or after injecting carrageenan or CFA into the paw indicating that neither MSK1 nor MSK2 controlled mechanical sensitivity of the paw either in naïve or inflamed condition. (E) CFA injection into the paw significantly increased oedema measured as paw thickness both in WT and MSK1^-/-^ mice 2 days after the injection. The increase in the thickness was more pronounced in MSK1^-/-^ then in WT mice. (two-way ANOVA, Bonferroni post-hoc test, ** and *** indicate, respectively, p<0.001 and p<0.0001). (F) – (I) Representative images of macrophages in WT ((F) and (H)) and MSK1^-/-^ mice ((G) and (I)) skin before ((F) and (G)) and 2 days after CFA injection ((H) and (I)) indicating that CFA induced macrophage activation and accumulation in both genotypes. Scale bar indicates 50 μm. (J) Quantification of macrophage activation/accumulation showed no difference between the two genotypes. (two-way ANOVA, Bonferroni post-hoc test, ** and *** indicate, respectively, p<0.001 and p<0.0001).

In contrast to the pronounced impact of MSK1 depletion on inflammatory heat hypersensitivity, hypersensitivity to punctate mechanical stimuli in the von Frey test developed normally in all genotypes in both inflammatory pain models (Figure 1C, D). These findings indicated a highly specific causal involvement of MSK1 in the development and persistence of inflammatory heat hypersensitivity.

### No reduction of inflammation in MSK1^-/-^mice

MSK1 is expressed in immunocompetent cells and involved in inflammatory reactions (Elcombe et al., 2013; Hossain et al., 2017; Kaiser et al., 2007; MacKenzie et al., 2013; Qi et al., 2021; Sattarifard et al., 2023). To elucidate whether MSK1 depletion prevented heat hypersensitivity through the attenuation of inflammatory processes, we assessed signatures of inflammation, such as paw swelling and macrophage accumulation. CFA injection resulted in a significant increase in paw diameter in both genotypes, and the degree of paw swelling was even significantly more pronounced in MSK1^-/-^than control mice (Figure 1E). No significant difference in inflammatory macrophage density was observed between MSK1^-/-^and WT mice two days after CFA injection (Figure 1F-J). Together, these findings suggested that the protection from inflammatory heat hypersensitivity in MSK1^-/-^mice was unlikely due to anti-inflammatory effects.

### MSK1 predominantly expressed in peptidergic nociceptors

Since we anticipated that MSK1 primarily impacted on neuronal processes augmenting nociceptive processing, we first assessed the location of MSK1 in relevant cell types within the nociceptive system. MSK1 immunoreactivity (IR) was detectable in dorsal root ganglia (DRG), the spinal dorsal horn and pain-related brain regions of WT but not MSK1^-/-^mice (Figure 2; Supplementary Figure 2A-I, Supplementary Figure 3, Supplementary Figure 4).

**Figure 2.**
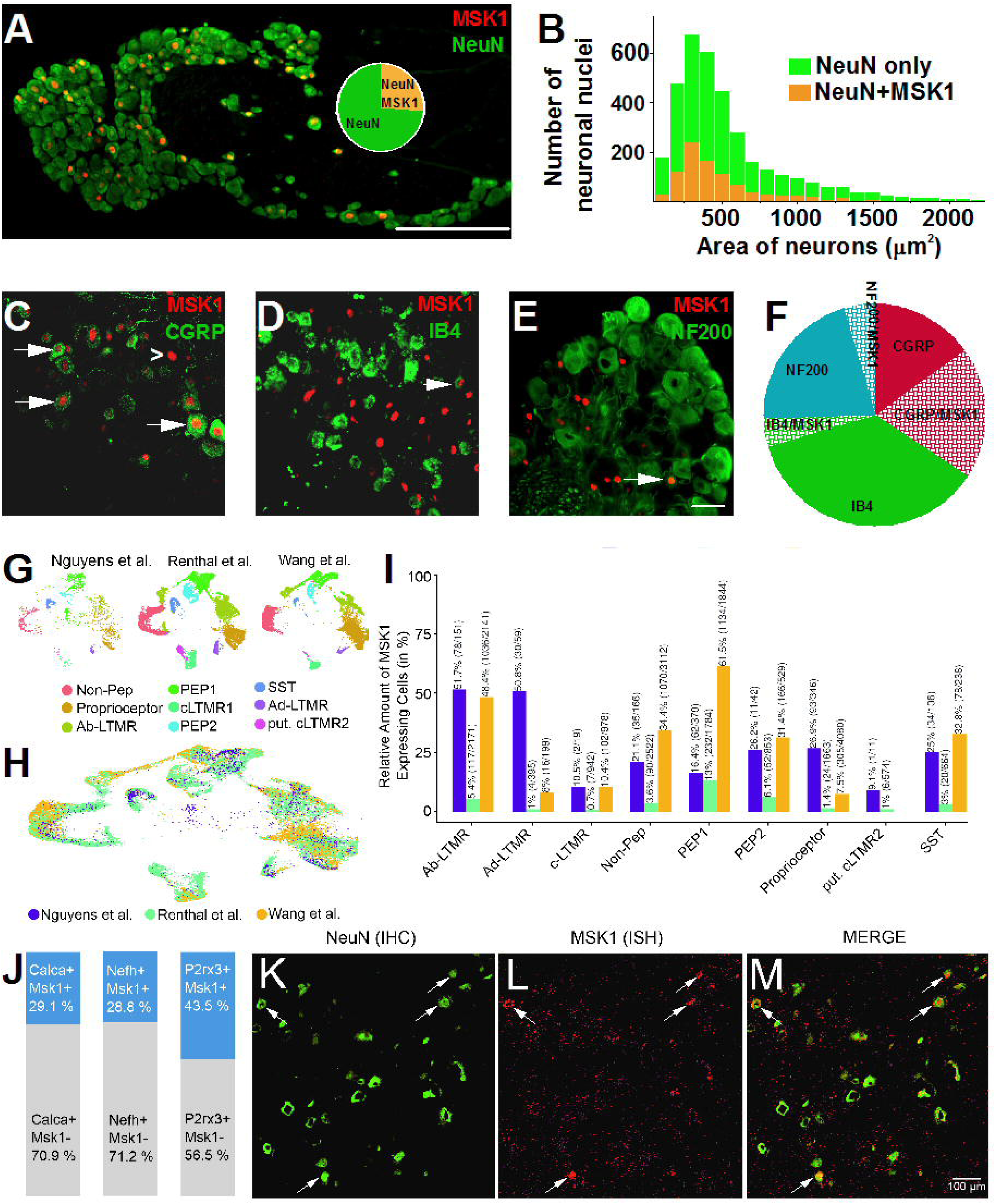
MSK1 is expressed in a group of mouse and human nociceptive primary sensory neurons. **(A)** MSK1 was expressed only in neuronal nuclei, identified by the specific marker NeuN, in mouse DRG. Scale bar indicates 250 μm. **(B)** The overwhelming majority of primary sensory neurons with nuclei exhibiting MSK1 expression had a small diameter. **(C)** – **(E)** MSK1 expression found mainly in CGRP-expressing primary sensory neurons (C). Only a few IB4-expressing (D) nociceptive neurons and NF200-expressing non-nociceptive neurons (E) exhibited MSK1 expression in the nucleus. Arrows indicate primary sensory neurons with MSK1-expression in the nucleus. Arrowhead in (C) indicates a CGRP-expressing neuron without MSK1 expression in the nucleus. Scale bar indicates 50 μm. **(F)** Proportion of the three major sub-types of primary sensory neurons with (hatched areas) or without (solid areas) MSK1 expression. **(G)** UMAPs showing clusters of human (snRNA-seq, purple) and mouse (snRNA-seq, turquoise; scRNA-seq, yellow) naive DRG neurons selected for MSK1-specific expression analysis. (H) UMAP showing integrated datasets shown in (G). (I) Bar chart depicts MSK1 expression in neuronal subtypes quantified in both relative and absolute terms in human (snRNA-seq, purple) and mouse (snRNA-seq, turquoise; scRNA-seq, yellow) naive DRG neurons. **(J)** Stack bars depict proportion of MSK1⁺ neurons exhibiting co-expression with *CALCA*, *NEFH*, and *P2RX3* in human DRG. (**K–M)** Microscopic images showing NeuN protein and *RPS6KA5* (MSK1) mRNA expression pattern in human DRG. Arrows indicate DRG neurons co-expressing *RPS6KA* mRNA and NeuN. Scale bar: 100 μm.

In the DRG, MSK1 was exclusively expressed in neurons, but not in non-neuronal cells (Figure 2A). Overall, 26±1.3 % (N=3 mice) of all neurons exhibited nuclear MSK1-IR (Table 1). The majority of MSK1-IR neurons were small diameter cells, which are generally accepted to possess nociceptive function (Nagy, 2004); Figure 2B). Among the three main distinct sub-populations of primary sensory neurons, more than 50 % of neuropeptide-containing (peptidergic; immunoreactive for example for the neuropeptide calcitonin gene-related peptide (CGRP)) nociceptors expressed MSK1 (Figure 2C, F; Table 1). Only about 10 % of the non-peptidergic (identified by isolectin B4 (IB4)-binding) nociceptors, and 18 % of non-nociceptive neurons (immunoreactive for heavy weight neurofilament (NF200)), were IR for MSK1 (Figure 2D, E and F; Table 1). In turn, three quarters of all MSK1-IR neurons in the DRG were peptidergic, 16 % were IB4-binding and 18 % NF200-IR cells (Figure 2F; Table 1).

**Table 1:** Number of neurons expressing MSK1 and markers of non-nociceptive and nociceptive primary sensory neurons in WT mice’ L3-L5 DRGs.

|  | **Total**<br>**absolute** | **Total**<br>**relative** | **n** |
| --- | --- | --- | --- |
| **Neuron** | 9003 | n.a. | 3 |
| **MSK1** | 2353 | 26.19±1.29 | 3 |
| **CGRP** | 3142 | 34.91±0.94 | 3 |
| **IB4** | 3704 | 40.4±3.88 | 3 |
| **NF200** | 2334 | 25.55±205 | 3 |
| **MSK1/CGRP** | 1777 | 19.8±0.5 | 3 |
| **MSK1/IB4** | 383 | 4.23±0.11 | 3 |
| **MSK1/NF200** | 411 | 4.58±0.19 | 3 |
| **MSK1 in CGRP** | 1777 | 56.86±2.66 | 3 |
| **CGRP in MSK1** |  | 76.34±3.43 | 3 |
| **MSK1 in IB4** | 383 | 10.62±0.73 | 3 |
| **IB4 in MSK1** |  | 16.32±0.75 | 3 |
| **MSK1 in NF200** | 411 | 18.33±2.34 | 3 |
| **NF200 in MSK1** |  | 17.68±0.81 | 3 |

n refers to the number of animals.

To explore whether MSK1 was also expressed in respective neuron populations in human DRG, we first employed an *in silico* approach, and reanalysed publicly available human single nucleus (sn) RNA-sequencing (RNAseq) datasets which include DRG from five patients (Nguyen et al., 2021). The percentage of neurons expressing *RPS6KA5* (the gene coding for MSK1) in human lumbar (L) 4-5 DRG ranged from 5.3% up to 33.3 % (Supplementary Figure 5). For cross-species comparison, the human dataset was integrated with single cell (sc) and sn RNAseq datasets from mice (Renthal et al., 2020; Wang et al., 2021), where neuronal subpopulations had been characterized based on transcriptomics signatures (Figure 2G). Non-nociceptive (Aβ-low threshold mechanoreceptors (LTMRs), Aδ-LTMRs, proprioceptive and three groups of nociceptive neurons (peptidergic (PEP1 and PEP2) and somatostatin (SST)-expressing) were particularly enriched with *RPS6KA5* and conserved between both species (Figure 2H and I). 29.1 %, 28.8 % and 43.5 % of *RPS6KA5* expressing nuclei, respectively, also expressed the gene encoding for CGRP (*CALCA)*, *NEFH* encoding for NF200 and the ionotropic purinergic receptor *P2RX3* which is characteristic for non-peptidergic nociceptors (Figure 2J). Using fluorescent *in situ* hybridisation (FISH), *RPS6KA5* mRNA was detected in human DRG in a subpopulation of neurons that were immunoreactive for the neuronal marker NeuN but also in compartments that were not clearly neurons (Figure 2K-M).

*MSK1 activation in nociceptive sensory neurons correlated with heat hypersensitivity* Based on its predominantly neuronal localisation, we hypothesised that MSK1 was activated in nociceptors upon tissue inflammation and quantified the number of DRG neurons expressing the activated form of MSK1 (p-MSK1) phosphorylated at Ser376 (Markou & Lazou, 2002). Very few p-MSK1 positive nuclei were detectable in L3-L5 DRGs collected from naive WT mice or one hour after saline injection, indicating very low MSK1 activity in sensory neurons innervating healthy tissue (Figure 3A and G; Supplementary Figure 6; Table 2). The relative number of p-MSK1 positive nuclei, and p-MSK1 expression were significantly increased at one hour and two days after CFA injection (Figure 3B, C, D and G; Table 2, Supplementary Figure 7). Recovery of MSK1 activation paralleled the reduction of heat hypersensitivity. Hence, when heat hypersensitivity recovered by ∼75 % thirty days after CFA injection, the proportion of p-MSK1-positive neurons was also reduced but still significantly higher than in naive mice (Figure 3E, F and G). Thus, the level of p-MSK1 indicating activation of the enzyme significantly correlated with heat hypersensitivity (Figure 3H). This suggested that the population of p-MSK1 expressing nociceptors could be causally and specifically involved in the development and persistence of inflammatory heat hypersensitivity.

**Figure 3.**
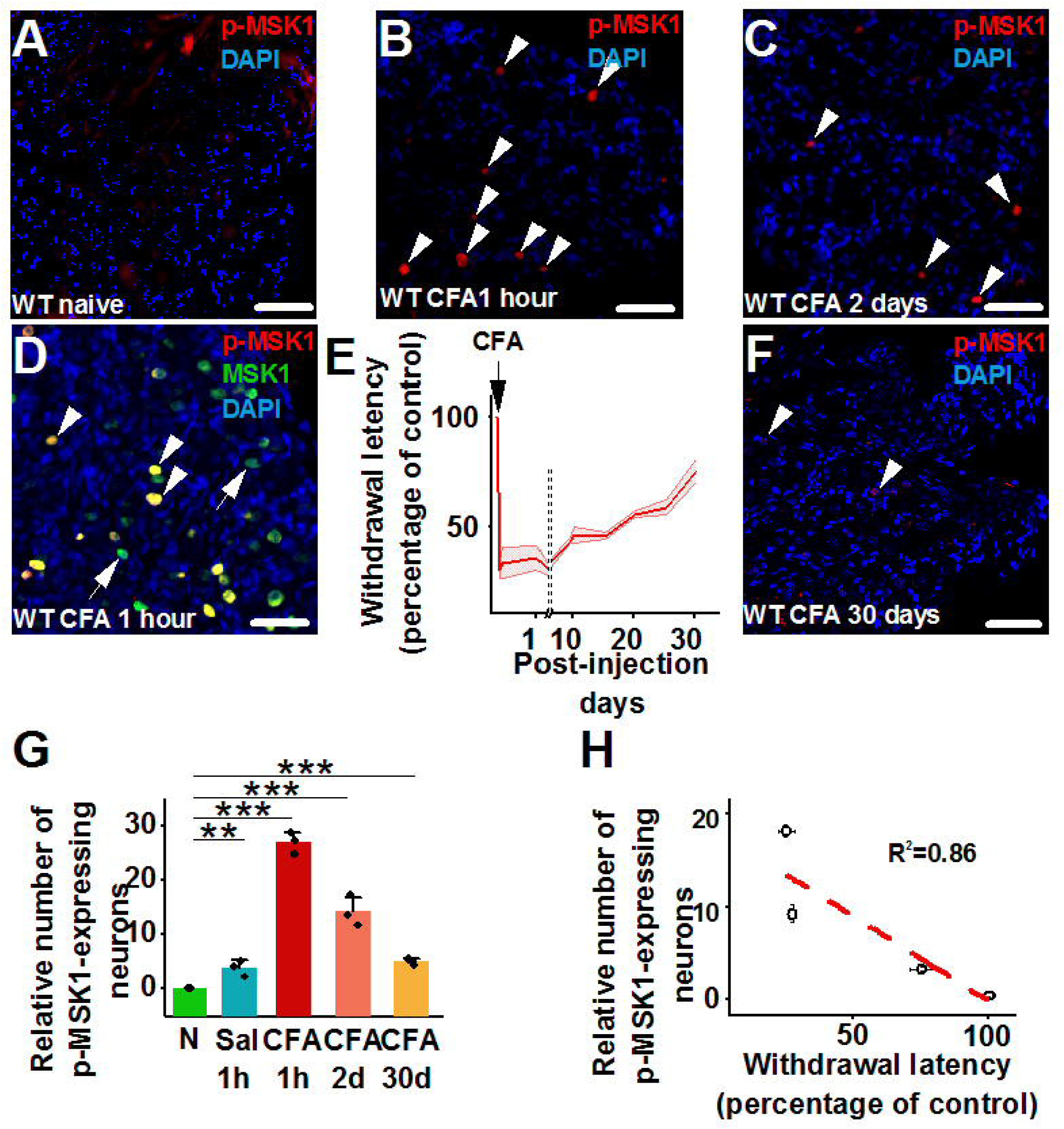
Inflammation induces MSK1 activation in a group of primary sensory neurons, which correlates with inflammatory heat hypersensitivity. **(A)** – **(C)** Microscopic images of WT DRG sections showing activated MSK1 (p-MSK1) expression in naive condition (A), and one hour (B), or two days after CFA injection into the hind paw. Arrowheads point to nuclei exhibiting p-MSK1 immunostaining. Scale bars indicate 50 μm. **(D)** Microscopic image of a WT DRG section showing MSK1 and p-MSK1 expression 1 hour after CFA injection into the hind paw. Arrowheads point to nuclei exhibiting MSK1 and p-MSK1 co-immunostaining, whereas arrows point to nuclei expressing MSK1 only. Scale bar indicates 50 μm. **(E)** Changes in paw withdrawal latency of WT mice injected with CFA into one hind paw between 1 hour and 30 days post-injection. Shaded area indicates SEM (n=5). **(F)** Microscopic image of a WT DRG section showing p-MSK1 expression 30 days after CFA injection into the hind paw. Arrowheads point to nuclei exhibiting p-MSK1 immunostaining. Scale bar indicates 50 μm. **(G)** Bar chart depicting relative number of p-MSK1 expressing neurons in naïve, one hour after saline (Sal 1h), one hour after CFA (CFA 1h), two days after CFA (CFA 48d) and 30 days after CFA (CFA 30d), injection into the hind paw (Fischer’s exact test, ** and *** indicate p=0.0007 and p<0.0001respectively; n=3 at each condition). **(H)** The relative number of p-MSK1-expressing neurons and paw withdrawal latency exhibit correlation (R^2^=0.86). As saline injection did not lead to heat hypersensitivity, withdrawal latency and p-MSK1 expression of saline-injected animals were not included.

**Table 2:** Number of neurons expressing p-MSK1 in naive condition and saline or CFA injection into the hind paw in WT mice.

|  | **Total**<br>**absolute** | **Total**<br>**relative** | **n** | **p** |
| --- | --- | --- | --- | --- |
| **naive** | 21 of 1751 | 0.7±0.36 | 6 | n.a. |
| **Saline, 1 hour** | 12 of 371 | 3.23±0.23 | 3 | 0.01 |
| **CFA, 1 hour** | 306 of 1119 | 27.34±1.01 | 3 | <0.0001 |
| **CFA, 2 days** | 221 of 1587 | 13.91±1.36 | 3 | <0.0001 |
| **CFA, 30 days** | 23 of 490 | 4.95±0.36 | 3 | 0.03 |

Statistical analysis was performed with Fisher's Exact test. n refers to the number of animals.

### p-MSK1 co-expressed with activated MAPK and p-CREB

MSK1 is directly activated by the MAPK extracellular signal-regulated kinase 1 and 2 (ERK1/2) and p38 (Deak et al., 1998; Drobic et al., 2010). Significantly more p-MSK1-positive nociceptors expressed p-p38 than p-ERK1/2 (Supplementary Figure 8A-D; Table 3). In agreement with MAPK involvement in inflammatory heat hypersensitivity (Ji et al., 2002), this points towards p38 as the main activator of MSK1 in nociceptors innervating inflamed tissues.

**Table 3:** Number of neurons expressing p-p38 or p-ERK1/2 with p-MSK and TRPV 1 2 days after CFA injection into the hind paw in WT mice.

| Marker | p-p38 |  |  | p-ERK1/2 |  |  |
| --- | --- | --- | --- | --- | --- | --- |
|  | Absolute number | Relative number | n | Absolute number | Relative number | n |
| **Naive** |  |  |  |  |  |  |
| **Total** | 1600 | 100 | 3 | 1243 | 100 | 3 |
| **p-MAPK** | 192 | 12.2±0.84 | 3 | 40 | 3.2 | 3 |
| **TRPV1** | 369 | 24.7±0.35 | 3 | 322 | 25.8±0.63 | 3 |
| **p-MAPK/TRPV1** | 87 | 5.6±0.55 | 3 | 23 | 1.9±0.5 | 3 |
| **CFA** |  |  |  |  |  |  |
| **Total** | 1132 | 100 | 3 | 1188 | 100 | 3 |
| **p-MAPK** | 340 | 29.9±0.48 | 3 | 126 | 10.6±0.48 | 3 |
| **TRPV1** | 421 | 37.2±0.24 | 3 | 416 | 35.2±0.71 | 3 |
| **p-MAPK/TRPV1** | 172 | 15.2±0.22 | 3 | 75 | 6.3±0.3 | 3 |
| **p-MSK1** | 183 | 16.2±0.54 | 3 | 190 | 16±0.5 | 3 |
| **p-MSK1/TRPV1** | 116 | 10.4±0.61 | 3 | 102 | 8.6±0.51 | 3 |
| **p-MAPK/p-MSK1** | 130 | 11.4±0.69 | 3 | 62 | 5.2±1.15 | 3 |
| **p-MAPK/p-MSK1/TRPV1** | 85 | 7.5±0.21 | 3 | 39 | 3.3±0.61 | 3 |
| **p (naive v CFA)\*** |  |  |  |  |  |  |
| **p-MAPK** | <0.0001 |  |  | <0.0001 |  |  |
| **TRPV1** | <0.0001 |  |  | <0.0001 |  |  |
| **p-MAPK/TRPV1** | <0.0001 |  |  | <0.0001 |  |  |

\*Fischer's Exact Test n refers to the number of animals.

MSK1 is also associated with the generation of p-CREB, which is increased in DRG neurons for example in experimental arthritis or burn injury (Segond von Banchet et al., 2016; Tamura et al., 2005; Torres-Perez et al., 2017). Importantly, the upregulation of p-CREB upon CFA injection was significantly reduced in MSK1^-/-^vs WT mice (Supplementary Figure 8E-H), indicating a major role of MSK1 regulating p-CREB in nociceptors during peripheral inflammation.

### MSK1 in nociceptors essential for inflammatory heat hypersensitivity

To document the importance of MSK1 in nociceptors for the development of inflammatory heat hyperalgesia further, we specifically down-regulated *Rps6ka5* in primary sensory afferents innervating the hind paw. Injection of an adeno-associated viral (AAV) vector carrying short hairpin (sh) RNA targeting *Rps6ka5* (AAV-*Rps6ka5*-shRNA) reduced MSK1 expression in DRG neurons compared to mice injected with AAV-scrambled shRNA (Supplementary Figure 9). Importantly, the mice that received AAV-*Rps6ka5*-shRNA developed significantly less heat hypersensitivity compared to mice injected with control AAV-scrambled shRNA (Figure 4A and B). This finding supports a critically important role of MSK1 in nociceptors for inflammatory heat pain.

**Figure 4.**
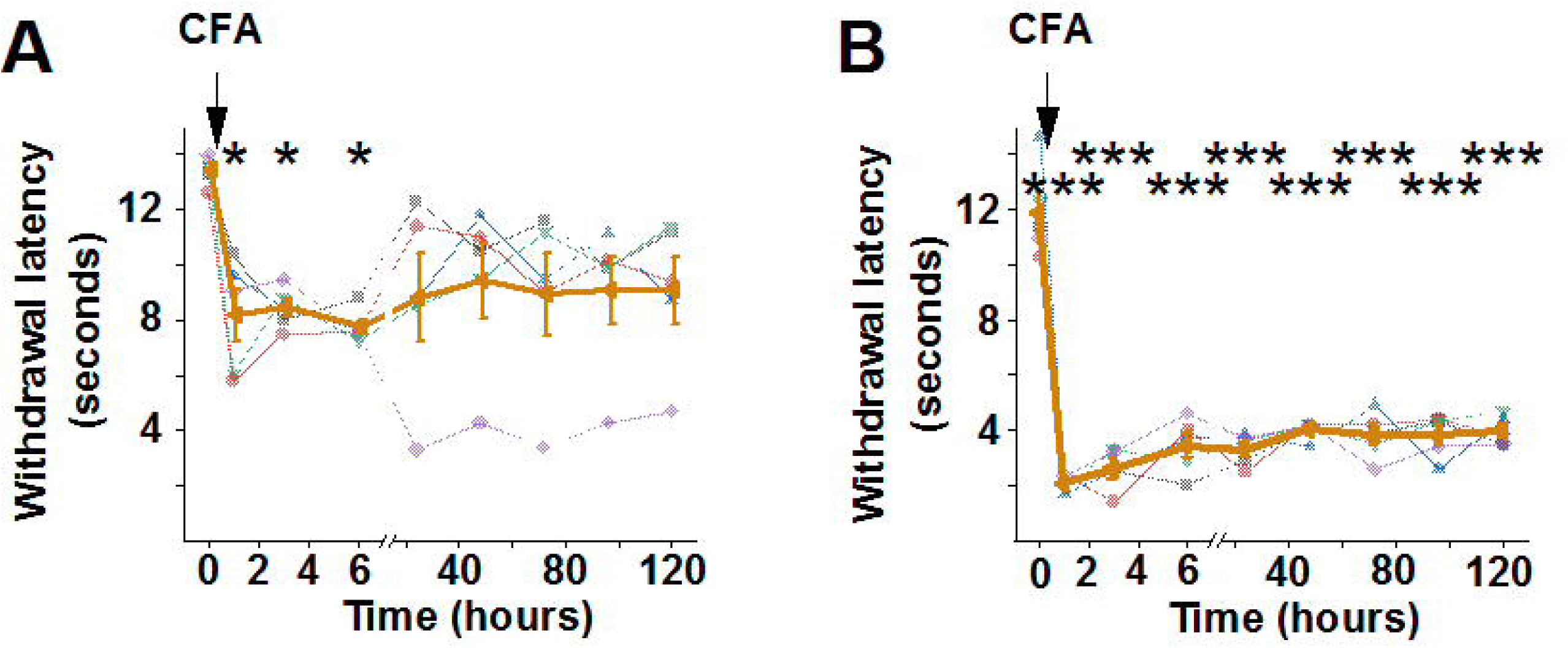
MSK1 in a group of primary sensory neurons is critical for inflammatory heat hyperalgesia **(A)** and **(B)** Paw withdrawal latency before and after CFA injection into the paw of mice that had been injected with AAV-*Rps6ka5*-shRNA (A) or AAV-scrambled shRNA (B) into the sciatic nerve 30-45 days before the assessment. Injection of AAV-*Rps6ka5*-shRNA attenuated the development and prevented the persistence of (A), injection of AAV-scrambled shRNA did not have any effect on inflammatory heat hypersensitivity (B); (two-way repeated measures ANOVA, Bonferroni post-hoc test, * indicate p=0.014, while *** indicate p<0.0001; n=5).

### TRPV1 expression regulated by MSK1 in nociceptors

MSK1 regulates the expression of a series of genes associated with adaptive responses (Choi et al., 2017; McGuire et al., 2017; Terazawa et al., 2020; Wagley et al., 2017). Based on the pivotal role of both TRPV1 and MSK1 in nociceptors in inflammatory heat hypersensitivity (Caterina et al., 2000; Davis et al., 2000) we hypothesised that MSK1 could regulate the expression of TRPV1 in nociceptors co-expressing *RPS6KA5* and *TRPV1*. Re-analysis of sc and sn RNAseq data revealed that a variable proportion of *RPS6KA5*+ cells co-express *TRPV1* (*RPS6KA5*+/*TRPV1*+) and the PEP1 peptidergic cluster exhibits the highest number of *RPS6KA5*+/*TRPV1*+-co-expressing neurons both in human (Nguyen et al., 2021) and mice (Renthal et al., 2020; Wang et al., 2021)(Figure 5A, B).

**Figure 5.**
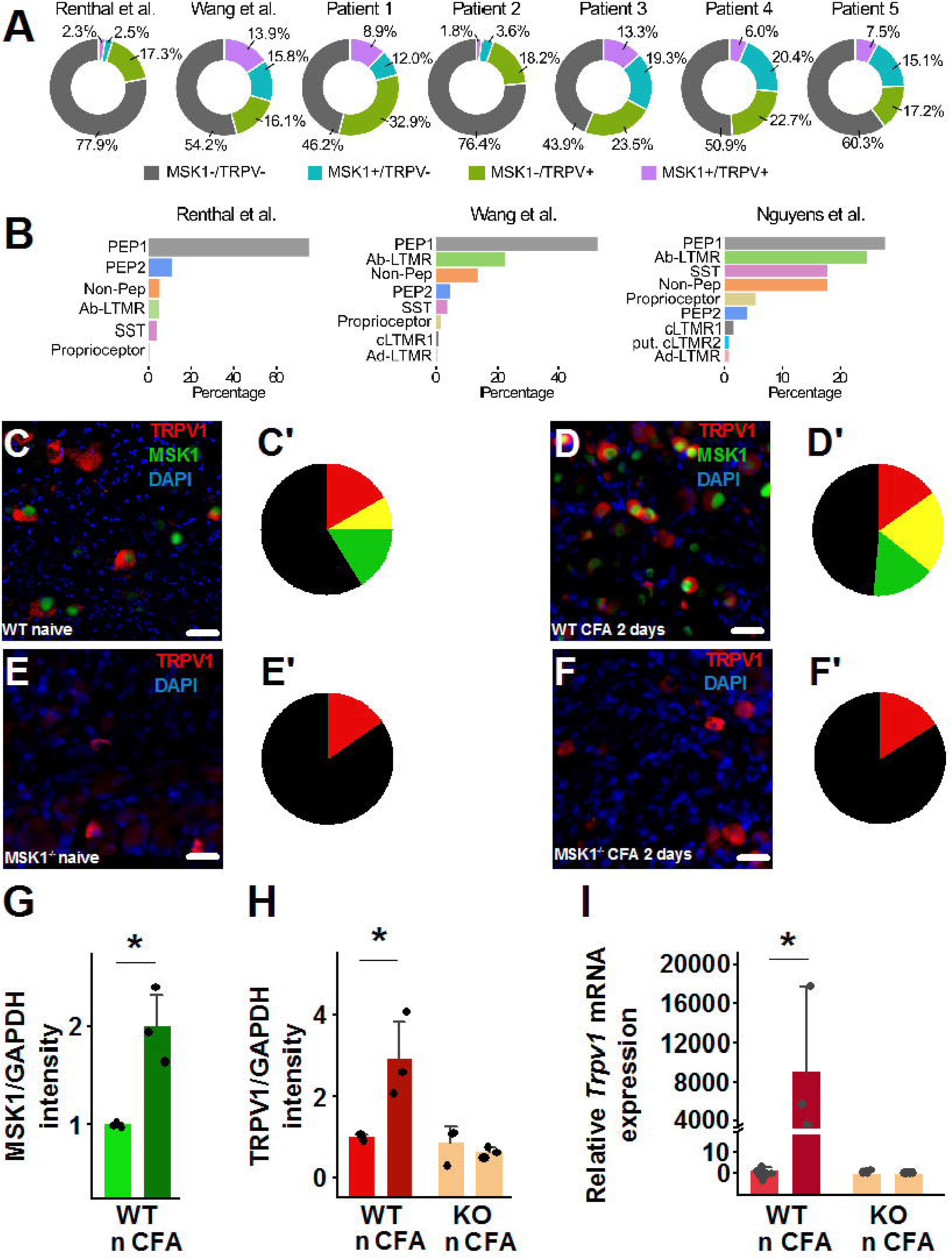
A group of primary sensory neurons co-expressing MSK1 and TRPV1 and MSK1 regulating TRPV1 expression. **(A)** Pie charts depict the relative number of *RPS6KA5/Rps6ka5* (MSK1)-and/or *TRPV1/Trpv1*-expressing mouse (Renthal et al., 2020; Wang et al., 2021) and human (Nguyen et al., 2021) (Patient 1-5) primary sensory neuron nuclei in publicly accessible databases (GSE155622, GSE154659, GSE168243). **(B)** Bar charts depict the relative number of *RPS6KA5/Rps6ka5* (MSK1)+/*TRPV1*/*Trpv1*+ primary sensory neuron nuclei in various transcriptionally-defined sub-classes of mouse (Renthal et al., 2020; Wang et al., 2021) and human (Nguyen et al., 2021) neurons. **(C)** and **(D)** Microscopic images show MSK1 and TRPV1 expression and co-expression in naive (WT naive) condition (C) and two days after CFA injection into the hind paw of WT mice (WT CFA 2days; D). Scale bars indicate 25 μm. **(C’)** and **(D’)** Pie charts depict the average relative number of MSK1 (green), TRPV1 (red) and MSK1/TRPV1 (yellow) expressing neurons. Inflammation significantly increases the relative number of MSK1/TRPV1-expressing neurons (Fischer’s exact test, p<0.0001 for all, n=3). However, the proportion of the TRPV1 only expressing cells (red) is not changed (Fischer’s exact test, p=0.184, n=3). **(E)** and **(F)** Microscopic images showing TRPV1 expression in naive condition (MSK1^-/-^ naive; E) and 2 days after CFA injection into the hind paw of MSK1^-/-^mice (MSK1^-/-^CFA 2 days; F). Scale bars indicate 25 μm. **(E’)** and **(F’)** Pie charts show the average relative number of TRPV1 (red) expressing neurons. Inflammation did not affect the relative number of TRPV1-expressing neurons in MSK1-/-mice (Fischer’s exact test, p=0.629, n=3). **(G)** Bar chart depicts relative MSK1 expression in WT mouse DRG in naïve condition (n) and 2 days after CFA injection (CFA) into one of the hind paws (Student’s t-test, * indicates p= 0.027 n=3). **(H)** Bar chart depicts relative TRPV1 expression in WT and MSK1^-/-^(KO) mouse DRG in naive condition (n) and 2 days after CFA injection (CFA) into one of the hind paws (two-way repeated measures ANOVA, Bonferroni post-hoc test, * indicate p=0.009 n=3). **(I)** Bar chart depicts *Trpv1* expression in WT and MSK1-/-(KO) mouse DRG in naïve condition (n) and 2 days after injecting CFA into one of the hind paws (CFA) by real-time quantitative PCR (ANOVA, Bonferroni post-hoc test, * indicate p<0.0005).

In naive mice only a small fraction of the entire neuron population in the DRG expressed both MSK1-IR and TRPV1-IR (Figure 5C and C’; Table 4) and the proportion of TRPV1-IR DRG neurons was significantly lower in naive MSK1^-/-^mice (Figure 5C-F’; Table 4) indicating a possible role of MSK1 in the regulation of TRPV1 expression. In WT mice, in line with our hypothesis, the percentage of TRPV1-IR or MSK1-IR neurons, the levels of TRPV1 and MSK1 proteins and *Trpv1* mRNA significantly increased after CFA injection in L3-L5 DRG harbouring the cell bodies of primary afferents innervating the hind paw (Figure 5C-D’, G-I; Table 4; Supplementary Figure 10). Most importantly, the percentage of neurons co-expressing MSK1-IR and TRPV1-IR, and also p-MSK1-IR and TRPV1-IR significantly increased after CFA injection in WT mice (Figure 5C-D’; Table 4; Table 5). Further, p-MSK1-, TRPV1-and MAPK-, particularly p-p38-IR were co-expressed in a significant number of neurons in WT mice (Supplementary Figure 11A and B). In DRG of MSK1^-/-^mice, *Trpv1* levels, the proportion of TRPV1-IR neurons as well as TRPV1 protein levels did not increase after CFA injection (Figure E-F’, H and I; Table 4; Supplementary Figure 10). Similarly, the CFA injection-induced TRPV1 upregulation in L3-5 DRG of WT mice whose sciatic nerve was injected with AAV-*Rps6ka5*-shRNA was significantly attenuated and was not different from the expression found in MSK1^-/-^mice’s DRG (Supplementary Figure 12).

**Table 4:** Number of neurons expressing MSK1 and/or TRPV1 in naive condition and 2days after CFA injection into the hind paw in WT and MSK-/- mice.

| **WT** | **Naive** |  |  | **CFA-injected** |  |  | **p** |
| --- | --- | --- | --- | --- | --- | --- | --- |
|  | **Total Absolute** | **Total Relative** | **n** | **Total Absolute** | **Total Relative** | **n** |  |
| **Neurons** | 1630 | n.a. | 3 | 1627 | n.a. | 3 | n.a. |
| **MSK1** | 403 | 24.8±0.34 | 3 | 586 | 36±0.82 | 3 | **<0.0001** |
| **TRPV1** | 412 | 25.3±0.36 | 3 | 581 | 35.7±0.76 | 3 | **<0.0001** |
| **MSK1/TRPV1** |  | 8.3±0.83 | 3 |  | 20.3±0.18 | 3 | **<0.0001** |
| **MSK1 in TRPV1** | 133 | 33.3±2.89 | 3 | 331 | 56.5±0.93 | 3 | **<0.0001** |
| **TRPV1 in MSK1** |  | 32.6±2.97 | 3 |  | 57±0.88 | 3 | **<0.0001** |
| **TRPV1 only** | 279 | 17.03±0.67 | 3 | 250 | 15.3±0.63 | 3 | **0.184** |
| **MSK1 only** | 270 | 16.51±0.53 | 3 | 255 | 15.6±0.69 | 3 | **0.505** |

| **MSK1<sup>-/-</sup>** | **Total Absolute** | **Total Relative** | **n** | **Total Absolute** | **Total Relative** | **n** | **p** |
| --- | --- | --- | --- | --- | --- | --- | --- |
| **Neurons** | 1620 | n.a. | 3 | 1582 | n.a. | 3 | n.a. |
| **TRPV1** | 252 | 15.4±1.09 | 3 | 256 | 16.1±0.61 | 3 | **0.629** |

Statistical analysis was performed with Fisher's Exact test
p (TRPV1 only in naive WT versus TRPV1 in naive MSK1<sup>-/-</sup>) =0.2358
p (TRPV1 only in inflamed WT versus TRPV1 in inflamed MSK1<sup>-/-</sup>) =0.5293
p (TRPV1 in naive WT versus naive MSK1<sup>-/-</sup>) <0.0001
p (TRPV1 in inflamed WT versus inflamed MSK1<sup>-/-</sup>) <0.0001
n refers to the number of animals.

**Table 5:** Number of neurons expressing p-MSK1 and/or TRPV1 in naive condition and 2days after CFA injection into the hind paw in WT mice.

|  | Naive |  |  | CFA-injected |  |  | p |
| --- | --- | --- | --- | --- | --- | --- | --- |
|  | Total absolute | Total relative | n | Total absolute | Total relative | n |  |
| Neuron | 1445 | n.a. | 3 | 1608 | n.a. | 3 |  |
| p-MSK1 | 21 | 1.42±0.26 | 3 | 242 | 15.08±0.41 | 3 | <0.0001 |
| TRPV1 | 373 | 25.98±0.95 | 3 | 591 | 36.71±1.03 | 3 | <0.0001 |
| p-MSK1/TRPV1 |  | 0 | 3 |  | 10.3±0.9 | 3 | <0.0001 |
| p-MSK1 in TRPV1 | 0 | 0 | 3 | 166 | 28.33±3.25 | 3 | <0.0001 |
| TRPV1 in p-MSK1 |  | 0 | 3 |  | 68.4±5.03 | 3 | <0.0001 |

Statistical analysis was performed with Fischer's Exact test. n refers to the number of animals.

Reanalysis of a publicly available RNAseq dataset (Pierrelée et al., 2021) of DRG transcriptome after carrageenan injection further supports MSK1/TRPV1 co-regulation as the expression of *Rps6ka5* and *Trpv1* transcripts increased after carrageenan injection (Supplementary Figure 13). The percentages of *Rps6ka5* expressing cells increased 48 hours after CFA injection (Renthal et al., 2020) with the strongest increase in PEP1 neurons (Supplementary Figure 13). These findings strongly support inflammation-associated transcriptional control of TRPV1 by MSK1 in a very specific population of nociceptors.

### TRPV1 functions fully segregated in DRG

We found that MSK1 expression defined two groups of TRPV1-expressing human and mouse DRG neurons (Figure 5). In CFA-injected MSK1^-/-^mice the proportion of neurons expressing TRPV1 only was similar to that of naive WT mice (Figure 5C’, D’, E’ and F’; Table 4) and depletion or downregulation of *Rps6ka5* prevented the development of inflammatory heat hypersensitivity without affecting acute baseline responses to noxious heat in healthy tissues (Figure 1; Supplementary Figure 1A; Figure 4). These findings together argue for a specific and critically important role of TRPV1 – MSK1 co-expressing neurons in inflammatory heat pain. This raises the possibility that this specific regulatory connection between MSK1 and TRPV1 was associated with a full functional segregation of TRPV1 and its role in acute versus inflammatory pain transduction by utilising distinct specific nociceptor lines.

To characterise the specific nociceptor line for the development and persistence of inflammatory heat hyperalgesia further, we combined indirect immunofluorescent staining and RNAscope labelling on L4-L5 DRG sections of naive mice for MSK1 co-expression with the three transient receptor potential (TRP) channels, TRP subfamily A member 1 (TRPA1), subfamily M member 3 (TRPM3) and TRPV1 participating in the transduction of noxious heat at the nociceptive nerve terminal (Figure 6A; (Vandewauw et al., 2018)). Among the 492 unequivocally identified neurons with clearly distinguishable nuclei, 62 (∼13 % of the total number of neurons) exhibited MSK1 expression together with *Trpv1* either alone or with other TRP channels, which was comparable with the proportion of the TRPV1-IR / MSK1-IR neurons (Figure 6A and B; Table 6). Of the 62 *Trpxx*+/*Rps6ka5*+ neurons, 18 expressed *Trpv1* only, which accounted for ∼3.6 % of the total number of neurons. This low proportion of the *Trpv1*+/*Rps6ka5*+ neurons in naive condition is in line with the hypothesis that TRPV1-expressing nociceptors are functionally segregated and inflammation-induced expansion of TRPV1-IR / MSK1-IR nociceptors are critical for inflammatory heat hyperalgesia (Figure 6A and B, Table 6).

**Figure 6.**
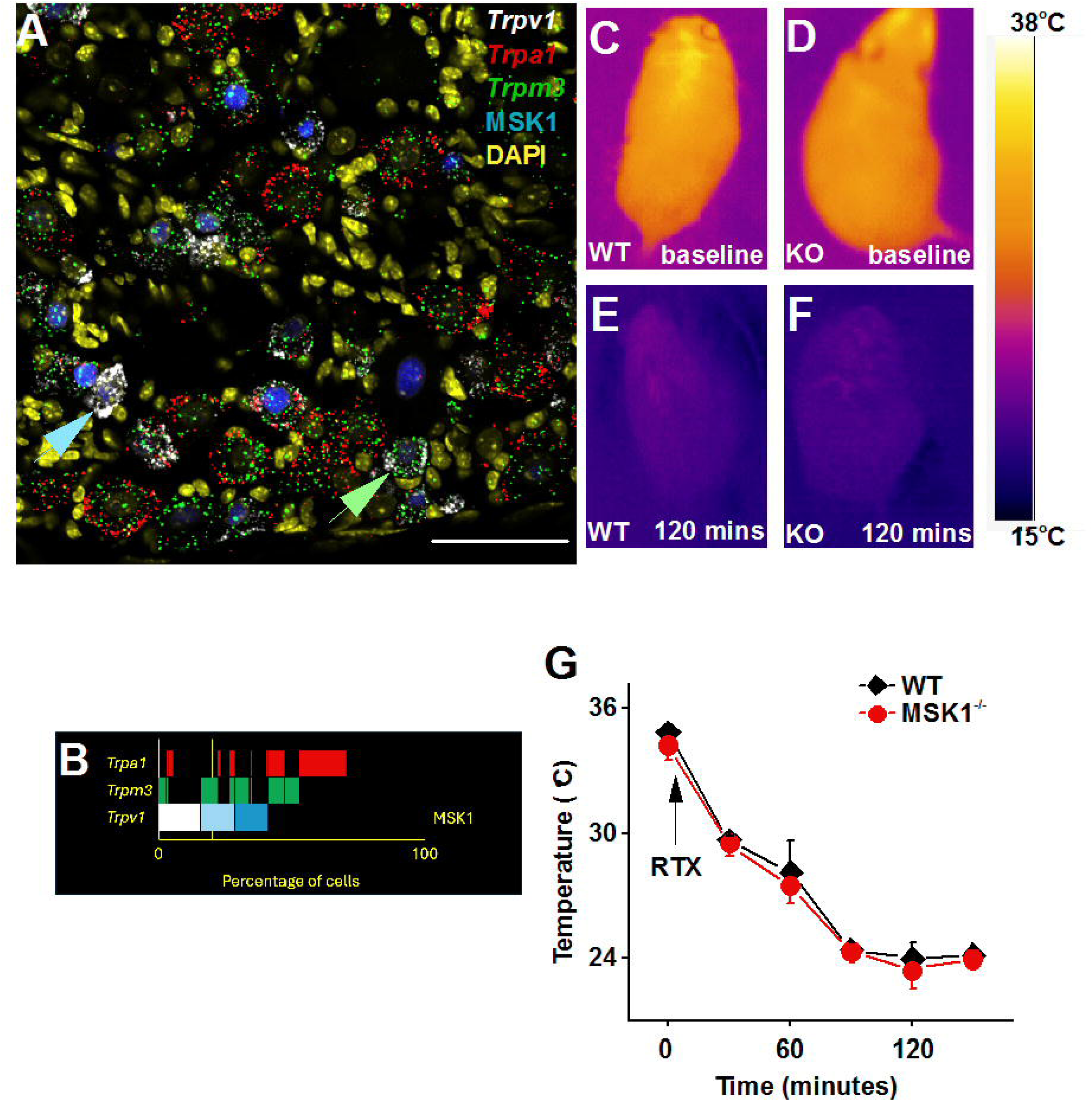
MSK1 expression defines a group of TRPV1-expressing nociceptors for inflammatory heat hyperalgesia **(A)** Microscopic image of a naive mouse DRG section showing *Trpm3*, *Trpv1*, *Trpa1* and MSK1 expression following combined labelling with triplex RNAscope (*Trpm3*, *Trpv1*, *Trpa1*) and immunostaining (MSK1). The image has been pseudo-coloured for visualisation. The blue arrow indicates a primary sensory neuron co-expressing *Trpv1* and MSK1 whereas the green arrow points to a neuron expressing *Trpv1* and *Trpm3* without MSK1. Scale bar indicates 50 μm. **(B)** Chart depicts quantified data of *Trpm3*/*Trpv1*/*Trpa1* triplex RNAscope labelling combined with MSK1 immunostaining. **(C)** – **(D)** Thermal images of WT (WT; C) and (E), and MSK1^-/-^ mice (KO; D) and (F) before (C) and (D), and 120 minutes after ip RTX injection (D) and (F). **(E)** Time course of changes in the highest surface temperature of WT (black diamond) and MSK1^-/-^mice (red dot). n=3. (Multiple repeated measures ANOVA, Bonferroni post-hoc test, p=0.42).

**Table 6:** Number of neurons expressing Trpv, Trpa1, Trpm3, and MSK in naive L3-L5 mouse dorsal root ganglia.

|  | Absolute number | Relative number |
| --- | --- | --- |
| Identified neurons with nucleus | 492 |  |
| *Trpv1* total | 133 | 27±0.27 |
| *Trpa1* total | 144 | 29.37±1.05 |
| *Trpm3* total | 138 | 28.13±1.25 |
| MSK1 total | 126 | 25.65±0.57 |
| *Trpv1* only | 55 | 11.1±0.68 |
| *Trpa1* only | 84 | 17.17±0.75 |
| *Trpm3* only | 27 | 5.56±0.63 |
| MSK1 only | 32 | 6.58±0.6 |
| *Trpv1/Trpa1* | 6 | 1.25±0.77 |
| *Trpv1/Trpm3* | 13 | 2.61±0.39 |
| *Trpa1/Trpm3* | 27 | 5.59±1.12 |
| MSK1/*Trpv1* | 18 | 3.62±0.55 |
| MSK1/*Trpa1* | 3 | 0.61±0.04 |
| MSK1/*Trpm3* | 26 | 5.3±1.2 |
| *Trpv1/Trpa1/Trpm3* | 2 | 0.39±0.2 |
| MSK1/*Trpv1/Trpa1* | 4 | 0.82±0.57 |
| MSK1/*Trpv1/Trpm3* | 32 | 6.47±1.04 |
| MSK1/*Trpa1/Trpm3* | 2 | 0.39±0.2 |
| MSK1/*Trpv1/Trpa1/Trpm3* | 8 | 1.62±0.17 |

To corroborate the hypotheses of a full segregation of TRPV1 functions further, we assessed the impact of MSK1 depletion on thermoregulation using systemic application of the ultrapotent TRPV1 agonist resiniferatoxin (RTX) to WT and MSK1^-/-^mice (Gentry et al., 2015). The baseline body temperature was similar in WT and MSK1^-/-^mice and, as expected, RTX injection resulted in a decreased body temperature in both genotypes excluding a regulatory effect of MSK1 on TRPV1 for thermoregulation (Figure 6C-G).

Together, these findings support the idea that the regulatory effect of MSK1 on TRPV1 expression indeed was specific for nociceptors mediating inflammatory heat hypersensitivity but not to acute heat pain and thermoregulatory processes.

### TRPV1+/RPS6KA5+ nociceptors’ function conserved in human

Based on our findings on the critically important role of MSK1 in inflammatory heat hyperalgesia (Figure 1 and 4) and the proposed regulation of TRPV1 expression in a group of nociceptors in mice (Figure 5), we anticipated that the newly discovered MSK1-TRPV1 interaction was conserved in human. Therefore, we assessed post-mortem human DRG from patient with and without history of the inflammatory disease arthritis (donor demographics in Supplementary Table 1). *TRPV1* as well as *RPS6KA5* expression were highly correlated and significantly upregulated in DRG obtained from arthritis patients as compared to controls without a painful disorder (Figure 7A -D).

**Figure 7.**
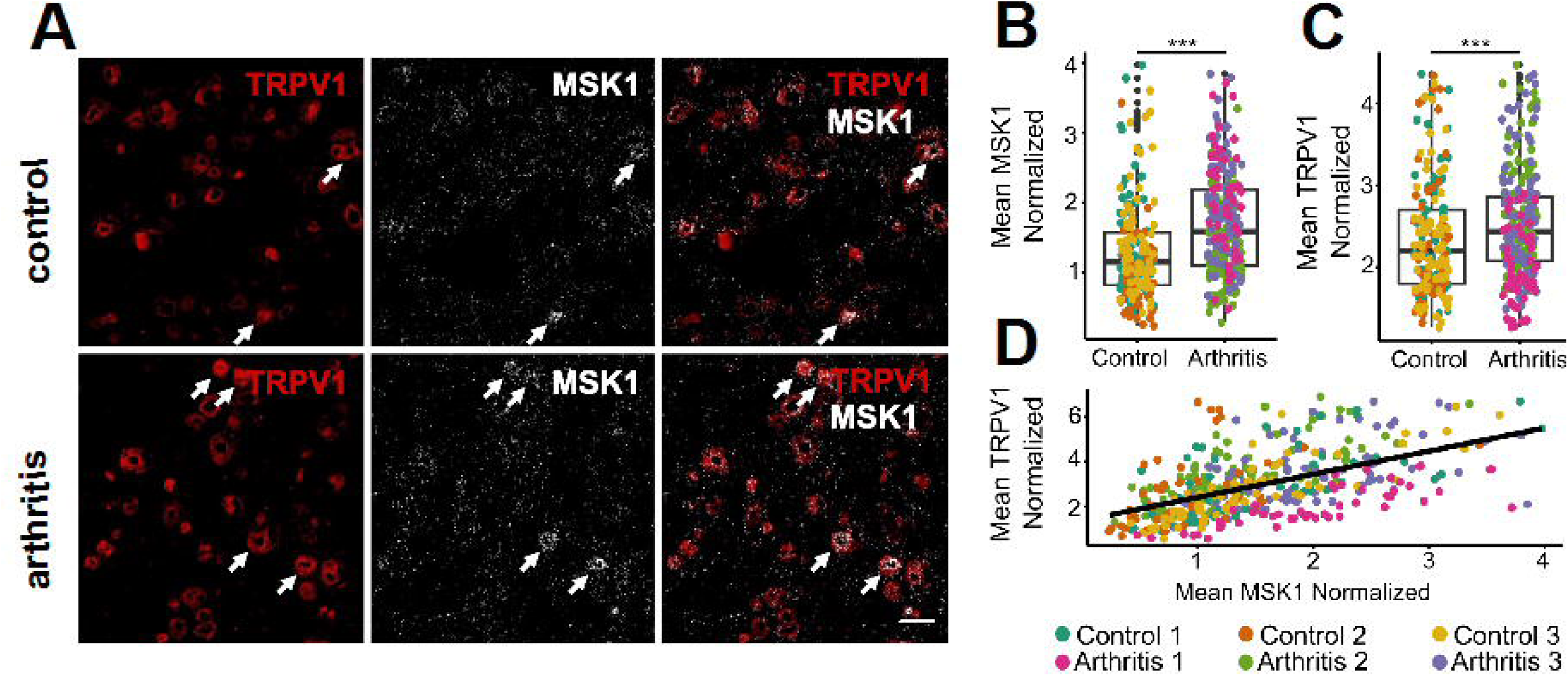
MSK1 and TRPV1 are co-regulated by arthritis in human primary sensory neurons **(A)** Microscopic images of human DRG sections from control and arthritic human donors showing TRPV1 and *RPS6KA5* (MSK1) expression using combined *in situ* hybridisation histochemistry and immunostaining. Arrows point to TRPV1 – *RPS6KA5*-co-expressing neurons. Scale bar indicates 100 μm. **(B)** – **(C)** depict *RP6KA5* (MSK1; H) and TRPV1 (I) expression levels and the correlation between *RP6KA5* (MSK1) and TRPV1 expression levels in primary sensory neurons of 3 control and 3 arthritis patients.

These findings, together with the significant attenuation of inflammatory heat hypersensitivity observed after silencing *Rps6ka5* in mouse nociceptors (see Figure 4A and B), support the idea that MSK1 is a major component of the regulatory machinery setting TRPV1 expression, and identifies a group of nociceptors essential for the development and persistence of inflammatory heat hyperalgesia.

## Discussion

Our results support a critically important role of MSK1 regulating TRPV1 expression in a specific group of MSK1-expressing nociceptors to initiate and maintain inflammatory heat hyperalgesia. We provide novel understanding of the pivotal role of TRPV1 in the development and persistence of inflammatory heat hypersensitivity, which is segregated from TRPV1’s other biological functions, such as acute noxious heat sensation and thermoregulation, both at cellular and regulatory levels (Vandewauw et al., 2018; Yue et al., 2022).

The essential role of TRPV1 in the development and persistence of burning inflammatory pain is generally well accepted (Caterina et al., 2000; Davis et al., 2000). Here, we found that MSK1 expression and activity are central for regulating TRPV1 to induce and maintain inflammatory heat hypersensitivity: global MSK1 depletion as well as MSK1 downregulation specifically in primary afferent sensory neurons prevented the inflammation-induced upregulation of TRPV1 expression and the development of inflammatory heat hypersensitivity. Tissue injury, even as small as injecting saline into the paw, and experimental inflammation activated and upregulated MSK1 in a specific subset of nociceptive primary afferents. The inflammation-induced p-MSK1 activity persisted for several weeks, and the number of neurons exhibiting p-MSK1 expression positively correlated with inflammatory heat hypersensitivity.

We propose a novel context-dependent “inflammation – MSK1 – *TRPV1* signalling cascade” involving inflammation-induced MSK1 upregulation, activation, and subsequent MSK1-dependent upregulation of TRPV1 expression in a specific population of primary afferent nociceptive neurons. Our data suggest that this signalling cascade is central and specific only for the development of inflammatory heat, but not mechanical hyperalgesia, since MSK1 depletion prevented the development of inflammatory heat but not mechanical hypersensitivity. Hence, we propose that MSK1 in TRPV1 expressing nociceptors serves as a specific checkpoint regulatory hub for the development and persistence of inflammatory heat hyperalgesia.

In contrast to pharmacological TRPV1 blockers used in the past, inhibiting TRPV1 expression by depleting MSK1 did not compromise the detection of acute noxious heat stimuli in healthy tissue nor thermoregulation (Chizh et al., 2007; Gavva, 2008). As a plausible explanation, MSK1 expression segregated two populations of TRPV1 expressing primary nociceptive neurons both in mouse and human DRG: the overall abundance of the nociceptor population expressing TRPV1 but not MSK1 was unaffected by experimental inflammation. In contrast, the second type of nociceptors expressing both TRPV1 and MSK1 was increased after experimental inflammation. These findings together with RNAseq data indicated that MSK1+/TRPV1+ sensory neurons constitute a specific nociceptor subtype that serves the initiation and maintenance of inflammatory heat hypersensitivity to protect of injured and inflamed tissues from further damage. The nociceptors co-expressing TRPV1 and MSK1 or *Trpv1* and *Rps6ka5* were sparse in naive murine and human. However, the percentage of MSK1+/TRPV+ nociceptors extended and exhibited pMSK1 expression in inflammatory condition suggesting that the inflammation-induced increase was due to *de novo* expression of the two signature proteins in this group of sensory neurons. Together, our findings introduce a novel nociceptor subtype co-expressing MSK1 with TRPV1 as the most relevant for the development and persistence of inflammatory heat hyperalgesia. This specific type of nociceptor is specifically affected by inflammatory processes and subject to plastic changes in inflammation, whereas the detection of noxious heat in healthy tissues and thermoregulatory functions is achieved by different TRPV1-expressing neurons that do not rely on MSK1 regulatory pathways.

The discovery of the segregation of the three TRPV1 functions both at the cellular and regulatory levels is of critical importance, because TRPV1 inhibitors to control inflammatory heat hyperalgesia so far have been disqualified as clinically applicable pain killers due to severe adverse effects (Szallasi & Sheta, 2012): they not only increase the detection thresholds for noxious heat in healthy tissues, with a considerable risk of severe burn injury, but also raise the body temperature and induce fever (Chizh et al., 2007; Gavva, 2008; Gavva et al., 2008; Szallasi & Sheta, 2012). In contrast, MSK inhibition alleviates inflammatory hypersensitivity but do not impact on acute pain detection or thermoregulation (current findings). Therefore, MSK1 – *Trpv1* signalling in the newly discovered inflammatory nociceptor type (“inflammatory heat nociceptors”) qualifies as specific labelled line for inflammatory heat hyperalgesia. Targeting MSK1 regulation of TRPV specifically in inflammatory nociceptors prompts an entire novel strategy with significant potential to effectively and specifically control inflammatory heat hyperalgesia without increasing the risk for burn injury or fever.

## Materials and methods

### Animals

All experiments on animals were carried out in accordance with the UK Animals (Scientific Procedures) Act 1986; European Communities Council Directive (86/609/EEC), guidelines of the Committee for Research and Ethical Issues of ISAP (Zimmermann, 1983), National Institutes of Health *Guide for the Care and Use of Laboratory Animals* (Revised Guidelines). ARRIVE guidelines were followed to report findings, and Good Lab Practice guideline was adhered in all experiments. The Animal Welfare and Ethical Review Body, Imperial College London, UK, the Animal Welfare Committee at Pécs University, the Animal Welfare Committee at the University of Debrecen and the National Scientific Ethical Committee on Animal Experimentation in Hungary approved all procedures which were conducted under a Home Office Project Licence.

We used C57BL6/J WT and MSK1^-/-^and MSK2^-/-^mice also with C57BL6/J background in the study. MSK1^-/-^and MSK2^-/-^mice were bred by crossing MSK1/2 double knock out mice (Arthur & Cohen, 2000; Wiggin et al., 2002) and C57BL6/J WT mice. Genotypes were confirmed with polymerase chain reaction. Mice were used, in approximately 50-50 % male and female ratio, at the age of 10-35 weeks (Irfan et al., 2025). Animals were randomly allocated to various studies. When feasible, samples were processed and analysed in a blinded fashion. Animals at all facilities were housed in a temperature-and humidity-controlled room with 12-hour light–dark and provided standard rodent chow and tap water *ad libitum*.

### Assessing pain-related behavioural and inducing tissue inflammation

Mice were acclimatised then trained on three consecutive days either with von Frey monofilaments (Bioseb, France) in a Perspex chamber placed on a plastic mesh flooring, or with the Hargreaves apparatus (Ugo Basile, Italy) in a Perspex chamber placed on glass flooring. On the fourth day, baseline sensitivity for mechanical or heat stimuli were determined after 60-75 minutes acclimatisation.

To assess sensitivity to mechanical stimuli, a series of successive calibrated von Frey monofilaments (0.008 – 2.0 g) were applied to the paw according to the up-down protocol and the withdrawal threshold recorded (Bonin et al., 2014; Deuis et al., 2017). For determining heat sensitivity, the infrared heat source of the Hargreaves apparatus was placed underneath the plantar surfaces of one of the hind paws and the withdrawal latency recorded.

Inflammation was induced under isoflurane anaesthesia by injecting 25 υl carrageenan, CFA, or saline subcutaneously into the plantar surface of the left hind paw. The paw withdrawal threshold or the paw withdrawal latency was then determined repeatedly. After carrageenan or saline injection, the assessments were performed at one, two and three hours, post-injection. After CFA injection, the assessments were done at two, four and six hours after the injection, then daily either up to 5 days or at every second or third day up to 30 days.

### Sciatic nerve injection

Under isoflurane anaesthesia, the sciatic nerves on both sides were exposed at the mid-thigh. 2 µl of AAV5-*Rps6ka5*-shRNA or AAV5-scrambled shRNA (Vector Biolabs) were injected into each side of the sciatic nerve with a 10 µl Hamilton syringe and Hamilton needle (NDL small RN ga34/15 mm/pst45°) and then muscle and skin wounds were closed by suturing. Gene knockdown was tested 4 weeks after AAV injection by Western blot. Behaviour was assessed 30 days after the sciatic nerve injection (Irfan et al., 2025).

### Tissue harvesting

Animals, when tissues were used for immunostaining or *in situ* hybridisation, were terminally anaesthetised by intraperitoneal pentobarbital (Euthatal, UK) then transcardially perfused with saline followed by 4 % paraformaldehyde for up to 60 minutes. The L3-L5 DRGs, the brain and paw skin were cryoprotected in 30 % sucrose and used for immunostaining. For *Trpm3*/*Trpv1*/*Trpa1* triplex RNAscope ISH combined with MSK1 immunostaining, L3-L5 DRGs were place into 30 % sucrose in 4% paraformaldehyde for overnight at 4°C.

When tissues were used for Western blotting or qPCR, animals were killed by cervical translocation, the desired tissues dissected and, respectively, snap frozen on dry ice or placed into RNAlater until processing.

### Human tissues

Human DRG tissue was sourced from Anabios, Inc (San Diego, CA; Supplementary Table 1). For this study, L3 DRGs from 6 female consented organ donors were selected. After extraction, DRG were immediately flash frozen and stored in liquid nitrogen.

### Immunostaining

Tissues were cryoprotected in 30 % sucrose in PBS 1X, embedded in Tissue-Teck OCT compound (VWR), and cut into 20-25 µm or 50 μm section with a cryostat. While 50 μm sections were processed as floating sections, 20-25 μm sections were mounted on a series of 3-aminopro-pyl-triethoxysilane-coated glass slides, where at least ∼100 μm was kept between sections placed on the same slide to avoid analysing the same cell twice.

Sections were thoroughly washed four times for 10 minutes in PBS containing 0.1-0.25 % Triton X-100 (PBST). Unspecific epitopes were blocked for 1 hour at room temperature with 5 – 10 % normal serum diluted in PBST. The blocking solution was replaced with the primary antiserum diluted in PBST supplemented with 1 % normal serum for overnight at room temperature. Sections were then washed with PBST and incubated in secondary antibody for 1 – 2 hours at room temperature diluted in PBS, followed by a final wash in PBS four times. Sections were cover slipped with ProLong Diamond Antifade Mountant medium either with or without DAPI (Invitrogen) or incubated for 5 minutes at room temperature with DAPI (1:10000) then mounted with Mowiol. When necessary, slides were incubated for 30 seconds at room temperature with TrueBlack^®^ Lipofuscin Autofluorescence Quencher (Cat No. 92401, Cell Signaling) after incubation in the primary antibody. When incubated as free-floating, section were captured on 3-aminopro-pyl-triethoxysilane-coated glass slides following incubation in secondary antibodies. Antibodies used in the work are listed in Supplementary Table 2.

### In situ hybridisation

Human DRG sections, were fixed in 4 % PFA for 15 minutes at 4°C and then washed in PBS before dehydration with increasing concentration of ethanol at room temperature. Slides were then dried for 5 minutes at room temperature then incubated for 10 minutes in hydrogen peroxide at room temperature. The MSK1 probe (Hs-RPS6KA5-C1 Cat No. 1311271-C1, ACD) was then added and incubated for 2 hours at 40°C. The slides were then washed followed by the amplification steps. AMP1, AMP2 and AMP3 were subsequently incubated at 40°C respectively for 30 minutes, 30 minutes and 15 minutes with 10 minutes wash between each step. For the signal development step, HRP-C1 was added and incubated for 15 minutes at 40°C followed by 10 minutes washing step with wash buffer. Opal™ 650 dye (1:500, Cat No. FP1496001KT, Akoya Biosciences) was incubated for 30 minutes at 40°C and then washed with wash buffer for 10 minutes. The HRP Blocker was finally incubated for 15 minutes at 40°C and washed with wash buffer for 10 minutes. The slides further underwent immunofluorescence staining as described above.

For *Trpm3*/*Trpv1*/*Trpa1* triplex RNAscope combined with MSK1 immunostaining, mouse L4-L5 DRGs, following cryoprotection in 30 % sucrose in 4 % paraformaldehyde, were embedded in Tissue Freezing Medium (Leica), and cut at 10 μm with a cryostat. Sections were treated with H_2_O_2_ followed by Protease III (RNAscope Multiplex Fluorescent v2 Assay, Cat No. 323110; ACD) for 30 minutes at 40°C, then hybridized with probes specific to mouse *Trpm3* (ACD, Cat. No. 459911), *Trpv1* (ACD, Cat. No. 313331-C2), and *Trpa1* (ACD, Cat. No. 400211-C3). Subsequently, sections were incubated overnight in anti-MSK1 primary antibody (1:500, Cell Signaling, C27B2 rabbit mAb #3489) with 2 % normal serum. Sections were then washed four times for 10 minutes with PBST (PBS containing 0.1 % Tween 20) and incubated in Alexa Fluor 594 goat anti-rabbit secondary antibody (1:500, Thermo Fisher Scientific, Cat. No. A11037) for 1 hour at room temperature diluted in PBS with 2 % normal serum, followed by a final wash in PBST four times. Sections were counterstained with DAPI and mounted with ProLong Diamond Antifade Mountant (Thermo Fisher Scientific) for confocal imaging.

### Western blotting

Tissues were homogenised in a buffer containing a proteinase and phosphate inhibitor cocktail (Thermo Scientific, UK). The lysates were centrifuged at 4°C for 30 minutes then the protein content quantified (BioRad, UK). Equal amounts of proteins were loaded on gel (SDS-PAGE, Invitrogen, UK) and separated by electrophoresis. Proteins were then transferred to PVDF membranes, which were washed in tris-buffered saline containing 0.1 % Tween (TBS-T) (Santa Cruz Biotechnology) three times for five minutes then blocked with 5 % bovine serum albumin (Sigma) in TBS-T for one hour at room temperature. Membranes were then incubated overnight at 4°C on a shaker with the primary antibody (Supplementary Table 2). On the following day, the membranes were incubated with Horse Radish Peroxidase (HRP)-conjugated secondary antibody for one hour at room temperature on a shaker (Supplementary Table 2), followed by incubation with a loading control (GAPDH; EDM Millipore Corp) for one hour at room temperature. Membranes were scanned by GeneGnome XRQ (SynGene), and the blots were analysed using Image J. The intensity of the specific band was measured and normalised to the intensity of the loading control. To compare different gels, the same standard control (spinal cord) was loaded into each gel. The specific bands were first normalised to the loading control (spinal cord), then to GAPDH (which is also normalised to the GAPDH of the standard control).

### Image analysis

Sections were examined and images taken with one of the following microscopes; Olympus BX51 epifluorescent microscope with Olympus DP74 camera (Olympus, Tokyo, Japan), Widefield – HF3-Zeiss Axio Observer, controlled by Zen software (Zeiss), Olympus IX83 microscope with a ORCA-Fusion Digital CMOS camera C14440-20UP, LSM 710 confocal laser scanning microscope (Carl Zeiss, Jena, Germany). The camera settings at each microscopes were standardised for taking images. Images of two – three sections were captured from each slide. The Image J (Fiji) software package was used to analyse the intensity of the staining of neurons identified individually as region of interest (ROI) by marking the perimeter of the nucleus and the cytoplasm. Only cells with visible nuclei were included in the analysis. The staining intensity of each ROI was measured and used to determine a threshold of positive labelling (Sousa-Valente et al., 2017). Data from at least three animals were averaged, and statistical analyses were performed. For figures, the brightness and contrast were adjusted, if needed, using Fiji/ImageJ. To make fluorescent images shown in figures consistent, virtual colours were selected on some images.

### Real time quantitative polymerase chain reaction

The total RNA was isolated from the DRG samples using the Arcturus Picopure RNA Isolation Kit (12204-01, Applied Biosystems) following manufacturer’s instructions. Briefly, DRGs where lysed with the kit’s extraction buffer at 42°C for 30 minutes, RNA washed in pre-conditioned columns with corresponding washing buffers and RNA eluted in a final volume of 14 µL of the corresponding elution buffer. RNA concentrations and quality were checked (NanoDrop 2000; Thermo Scientific). cDNA libraries were generated using the SuperScript VILO cDNA Synthesis Kit (11754; Invitrogen; 10 minutes at 25°C, 60 minutes at 42°C, and 5 minutes at 85°C). cDNA quality was checked using NanoDrop 2000.

Concentration of cDNA samples were adjusted to 220 ng/µL with MilliQ water. Three housekeeping genes (*Actb*, *Gapdh* and *Rpll3*α) were used to normalise the data and assess relative expression changes for the different genes studied.

Relative qPCR assays and data analysis were performed similarly as in previous publications (Evans et al., 2021). Reactions were performed in triplicates using the SYBR Premix Ex Taq (RR420, Takara) and the StepOne Plus Real-Time PCR System (Applied Biosystems). PCR reactions were performed as follows: 5 minutes at 95°C for 5 minutes and then 40 cycles of 15 seconds at 95°C and 34 seconds at 56°C; and 15 seconds at 95°C. The melting curves were assessed by increasing temperatures in 0.5°C steps from 60 to 95°C.

### Bioinformatics

All analysis were performed in R (version 4.2.2) using Seurat Package (version 4.3.0). In each single cell/nucleus sequencing batch, raw counts were log-scaled using ScaleData. Human-mouse orthologs were identified using nichenetr (version 2.0.0). The data integration along the RNA-slot was prepared by SelectIntegrationFeatures and FindIntegrationAnchors. The integrated data were scaled, followed by principle component and umap dimensional reduction analysis and unsupervised clustering. In the RNAseq dataset (GSE 161764), the raw counts were variance stabilized using VST of DeSeq2 (1.41.5)

### Statistics

The Shapiro-Wilk tests confirmed normal distribution of data, and statistical analysis was performed using appropriate tests. The number of neurons expressing/co-expressing various markers was compared using the Fisher’s exact test. Statistical analysis was performed by Student’s t-test or various forms of analysis of variance (ANOVA) followed by the post-hoc Bonferroni test as appropriate. Data are expressed as average ± standard error mean. A difference was regarded significant at p<0.05.

## ACKNOWLEDGEMENTS

IN, MK, DZ and KD-P have been supported by a joint project grant from the Austrian Science Fund and National Research, Development and Innovation Office (Hungary); IN has been supported by British Journal of Anaesthesia/Royal College of Anaesthetists Project Grant; RMF has been supported by a PhD studentship from the Indonesian Endowment Fund; DA has been supported by a PhD studentship from the Kingdom of Saudi Arabia, Ministry of Defence, The General Directorate of Health Services, Academic Affair; JVPT has been supported by MICIU/AEI/10 (RYC2021–034012-I) and the “European Union NextGenerationEU/PRTR” program (13039/501100011033); IN and JVTP have also been supported by the *Conselleria de Educación, Cultura, Universidades y Empleo* from the *Generalitat Valenciana* with a *Subvencion a grupos de investigación emergentes* (grant CIGE/2024/73).

## DATA AVAILABILITY

Raw and processed data are available from the corresponding author on request.

