## Supplementary Figure for "Inflammatory burning pain depends on a specific line of nociceptors"

**Supplementary Figures**

**Supplementary Figure 1**

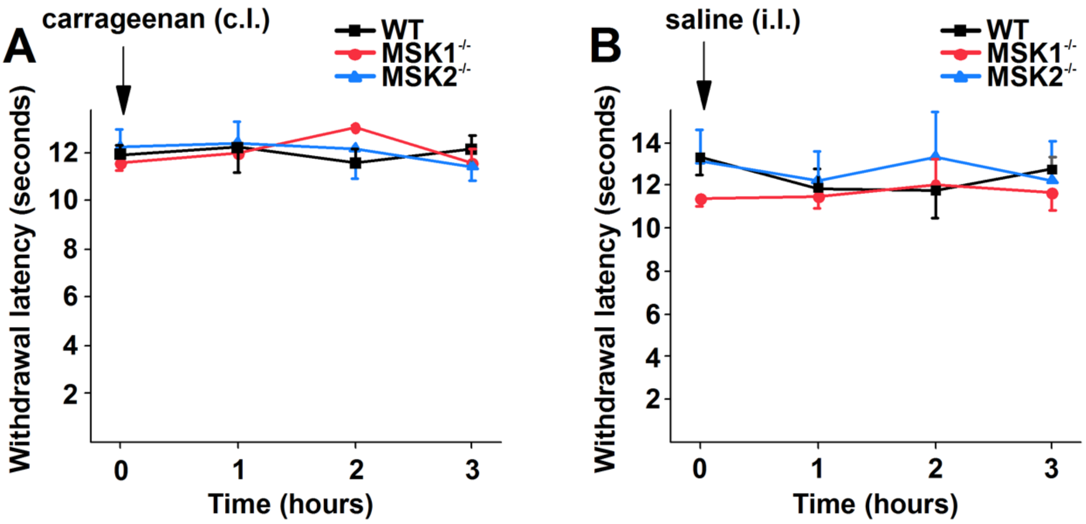


**Injecting the inflammatory agent carrageenan or saline does not induce hypersensitivity to heat stimuli, respectively on the contralateral or ipsilateral sides in WT, MSK1$^{-/-}$ or MSK2-/- mice.**

**(A)** and **(B)** Carrageenan (A) or saline (B) injection into the left paw does not reduce withdrawal latency, respectively, at the right or left paw in any genotype.

### Supplementary Figure 2

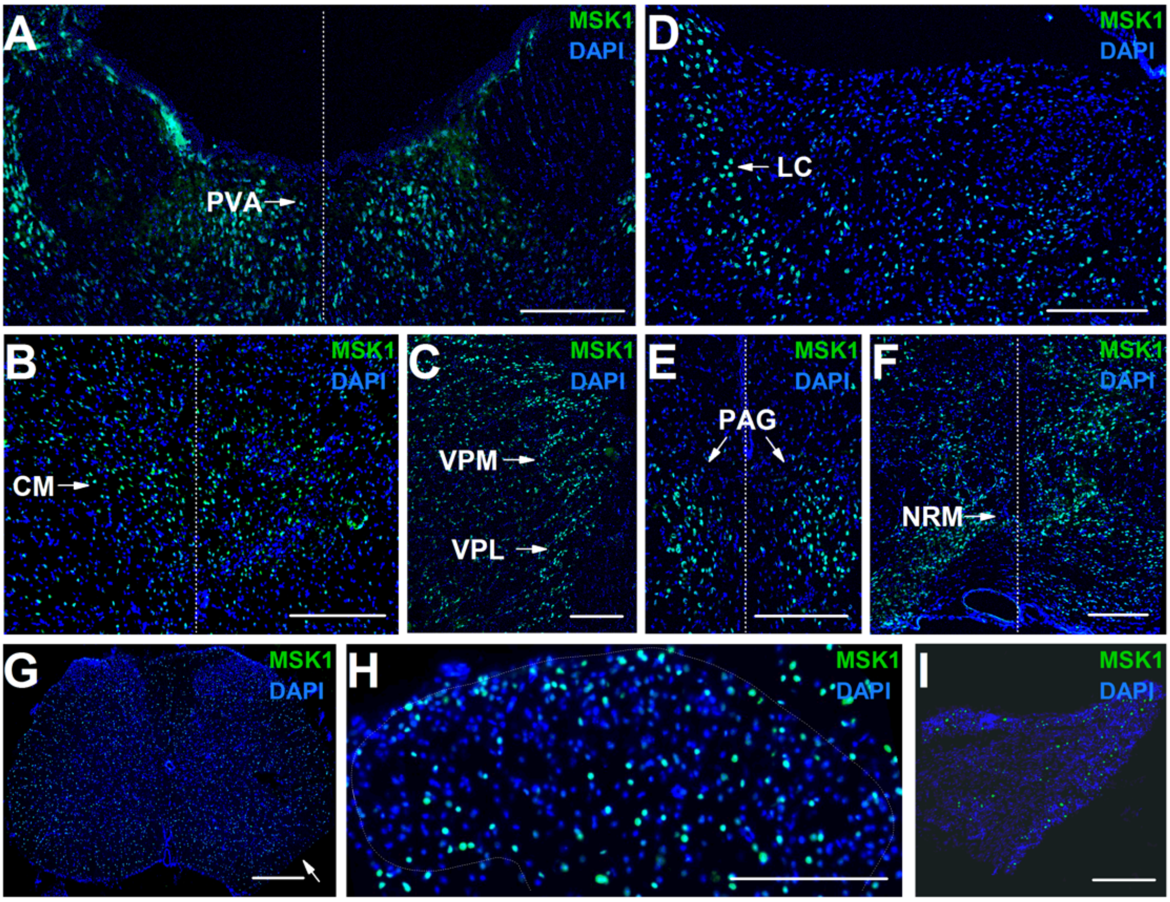


**MSK1 expressed in a series of nociception-related areas of the nervous system.**

**(A) – (I)** MSK1 expression or lack of expression in the paraventricular nucleus (PVA, (A)), centrum medianum (CM; B), ventral posteromedial (VPM) and ventral posterolateral (VPL) nuclei (C), locus coeruleus (LC, (D)), periaqueductal grey (PAG, (E)), nucleus raphe magnus (NRM; (F)), spinal cord (G) and (H) and dorsal root ganglion (I). Scale bars indicate 500 µm in (A) – (G) and 250µm in (H) – (I). Arrow in (G) indicates incision in the right ventral horn to identify the right side.

**Supplementary Figure 3**

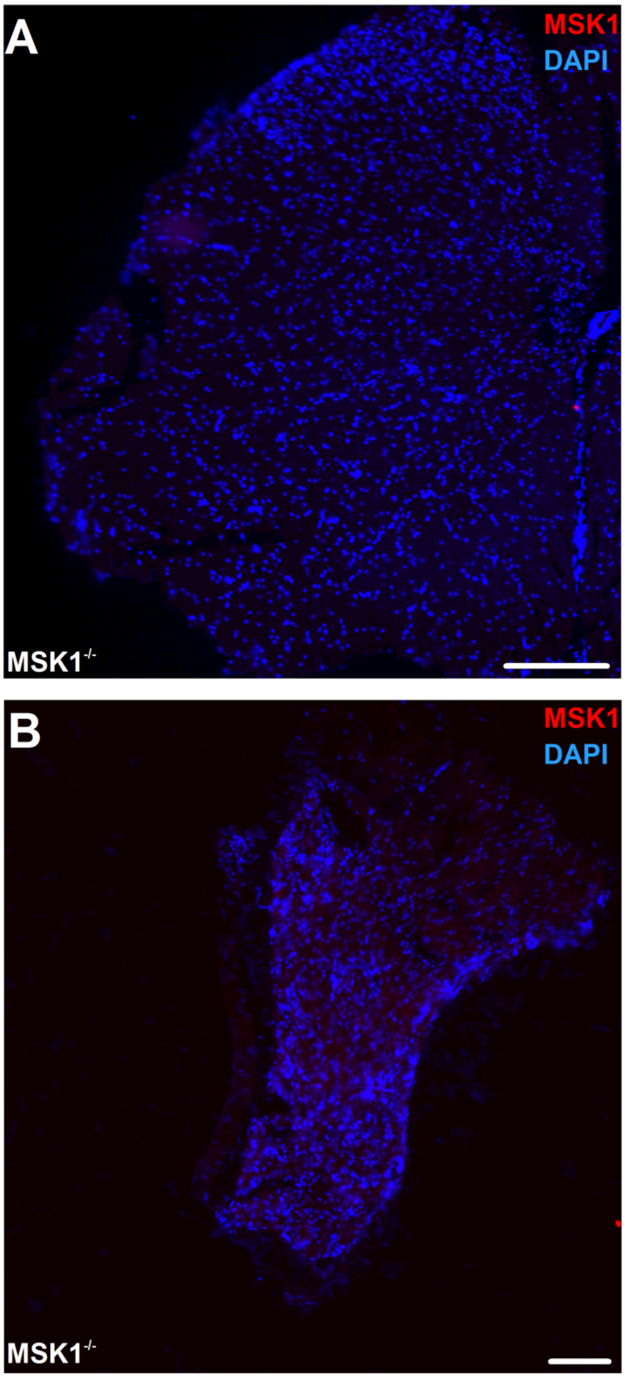


**The anti MSK1 antibody does not generate staining in the spinal cord or dorsal root ganglion collected from MSK1$^{-/-}$ mice.**

**(A)** and **(B)** Microscopic images of a spinal cord (A) and dorsal root ganglion section (B) cut from tissues collected an MSK1$^{-/-}$ mouse following the incubation in the anti-MSK1 antibody used throughout the study. No immunostaining is detectable in either the spinal cord (A) or dorsal root ganglion. Scale bars indicate 250 μm (A) and 100 μm (B).

**Supplementary Figure 4**

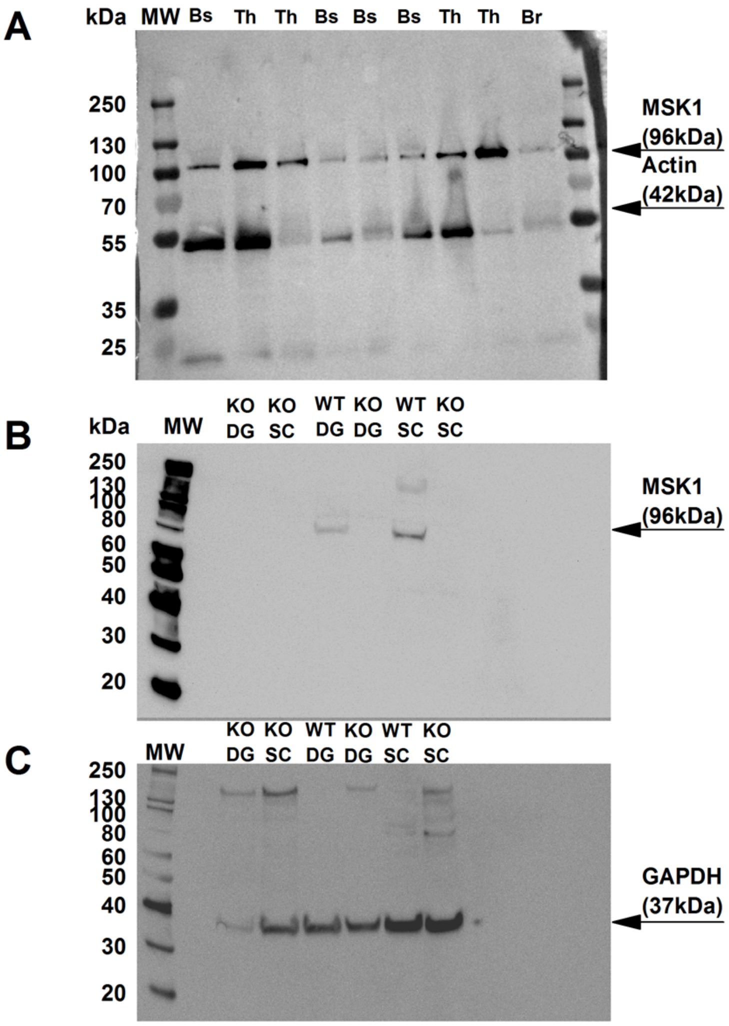


**The anti-MSK1 antibody specifically detects MSK1 in various parts of the nervous system.**

**(A)** – **(C)** Gel images of Western blotting of protein extracts of WT mouse brain stem (BS) and thalamus (Th; A) and WT and MSK1$^{-/-}$ (KO) mice's dorsal root ganglia (DG) and spinal cord (SC, B) probed with the anti-MSK1 antibody we used throughout the study (A and B), and the house keeping genes actin (A) or glyceraldehyde-3-phosphate dehydrogenase (GAPDH, C). The lane labelled with MW shows molecular weight markers.

**Supplementary Figure 5**

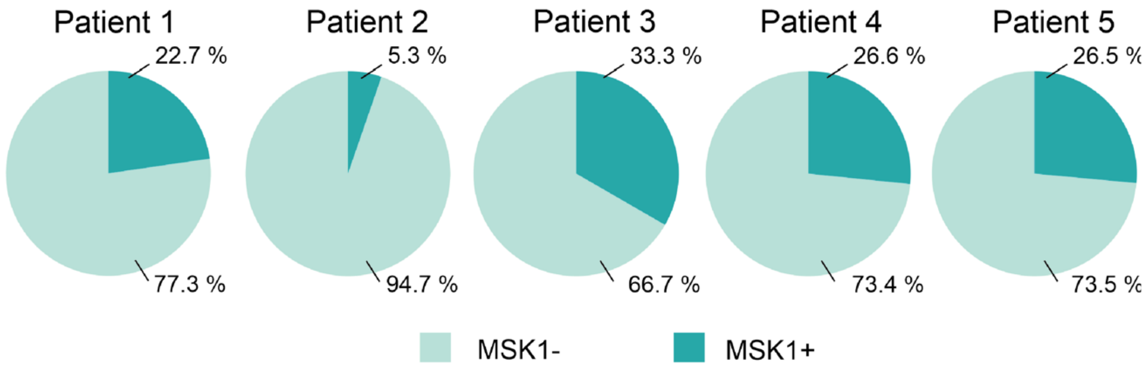


**MSK1 is expressed in a group of human primary sensory neurons**

Charts depict findings of reanalysing the human single nucleus (sn) RNA-sequencing (RNAseq) datasets of five patients (Nguyen et al. 2021) and show that *RPS6KA5* (mRNA transcribed from the gene coding for MSK1) is expressed in a group of primary sensory neurons.

**Supplementary Figure 6**

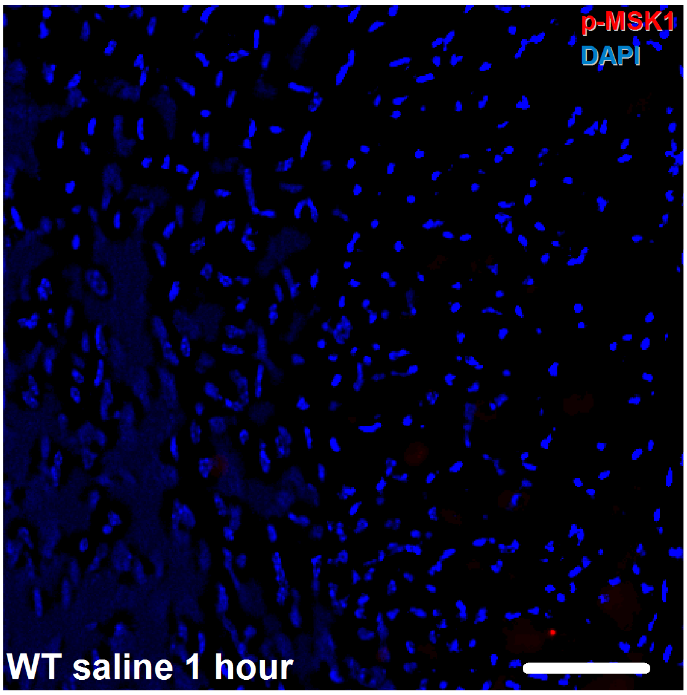


**Saline injection activates MSK1 in very few primary sensory neurons in WT mice.**

Microscopic image of section cut from a dorsal root ganglion collected from a WT mouse one hour after injecting saline into the hind paw. In this section, no activated MSK1 (p-MSK1)-expressing nucleus is visible. Scale bar indicates 50 μm.

**Supplementary Figure 7**

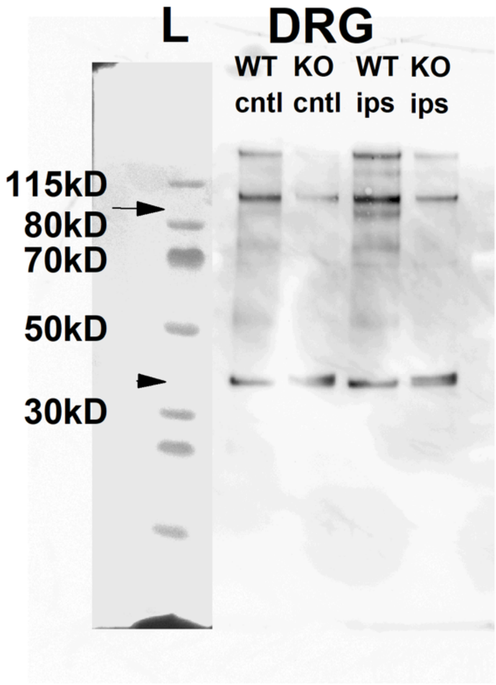


**Complete Freund's Adjuvant injection into the hind paw induces MSK1 activation in the ipsilateral L3-L5 dorsal root ganglion.**

Gel image of Western blotting of protein extracts of WT and MSK1$^{-/-}$ mice's (KO) L3-L5 dorsal root ganglia (DRG) and spinal cord (SC) collected from the ipsilateral (ips) and contralateral (cntl) side 2 days after CFA injection into the left hind paw. The lane labelled by L shows molecular weight markers. Arrow indicates the ~96kD, molecular weight of activated MSK1 (p-MSK1). Arrowhead indicates ~37kD, molecular weight of GAPDH.

**Supplementary figure 8**

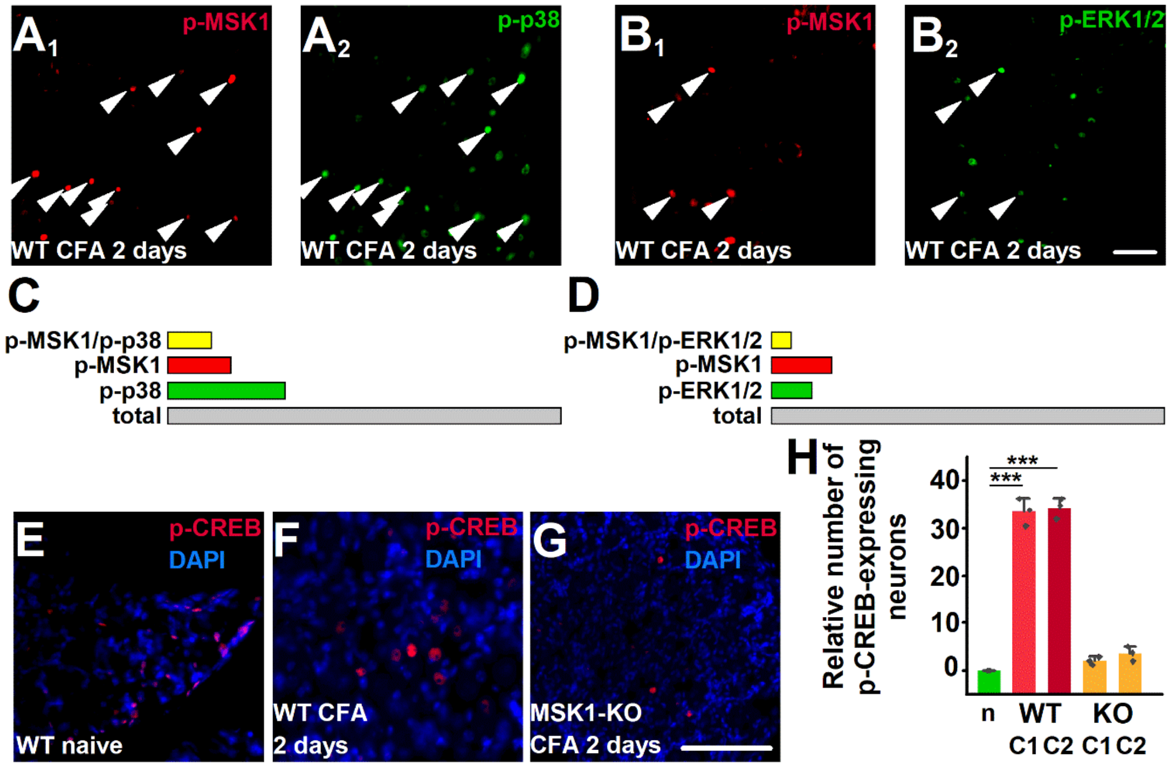


**Mitogen-activated protein kinases and cAMP response element binding protein act as up- and downstream signalling molecules, respectively, for MSK1.**

(A1 – B2) Microscopic images of WT DRG sections showing p-MSK1 (A1) and (B1) and p-p38
(A2) or p-ERK1/2 (B2) expression 2 days after CFA injection into the hind paw. Arrowheads point to nuclei exhibiting p-MSK1 and p-p38 or p-ERK1/2 co-immunostaining. Scale bar indicates 50 μm.
(C) and (D) Bar charts depicting the total relative number of p-MSK1, p-p38 (C) or p-ERK1/2 (D) and the relative number of neurons co-expressing p-MSK1 and p-p38 or p=MSK1 (C) and p-ERK1/2 (D) in WT mice's L3-L5 DRG 2 days after CFA injection into the paw.
(E – G) Microscopic images of WT (E – F) and (MSK1-/-) mouse L3-L5 DRG sections showing p-CREB expression in naive condition (E), and 2 days after CFA injection (F) and (G). In naive mice p-CREB is expressed in satellite cells. Scale bar indicates 100 μm.
(H) Bar chart depicting the relative number of p-CREB immunopositive neurons in WT and MSK1-/- (KO) mice DRG neurons in naive conditions, 1 day (C1) or 2 days (C2) after CFA injection (ANOVA followed by Bonferroni post-hoc test, *** indicate p<0.0001; n=3 at each condition).

### Supplementary Figure 9

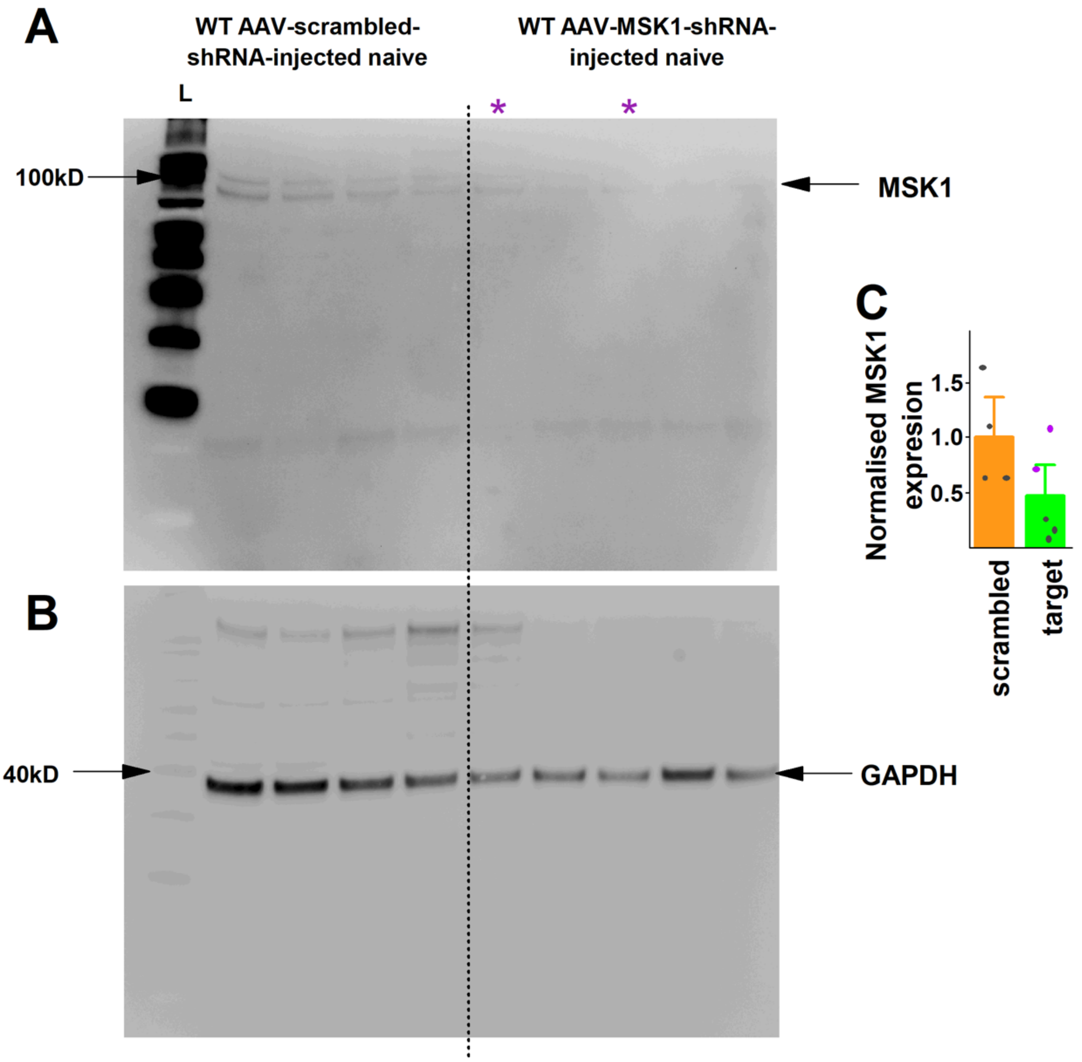


**Sciatic nerve injection of adeno-associated virus (AAV) serotype 5 vector carrying short hairpin (sh) RNA targeting *Rps6ka5* (AAV-*Rps6ka5*-shRNA) downregulates MSK1 expression in L3-L5 dorsal root ganglia in WT mice.**

**(A)** – **(B)** Gel images of Western blotting of protein extracts isolated from control WT mice's L3-L5 dorsal root ganglia (DRG) collected 30-45 days after injecting AAV-scrambled-shRNA or AAV-*Rps6ka5*-shRNA into the sciatic nerve show the effect of the two viral constructs on MSK1 expression (A). (B) shows GAPDH expression in the same samples. Vertical dotted line divides blots from scrambled and target shRNA-injected samples. Purple asterisks in AAV-*Rps6ka5*-shRMA indicate samples where downregulation appeared less successful.
**(C)** Chart depicts results of semiquantitative analysis of MSK1 expression in DRG from AAV-scrambled-shRNA and in AAV-*Rps6ka5*-shRNA-injected mice shown in (A) and (B). Purple dots indicate normalised MSK1 expression in samples labelled by purple asterisks on (A) and (B).

**Supplementary Figure 10**

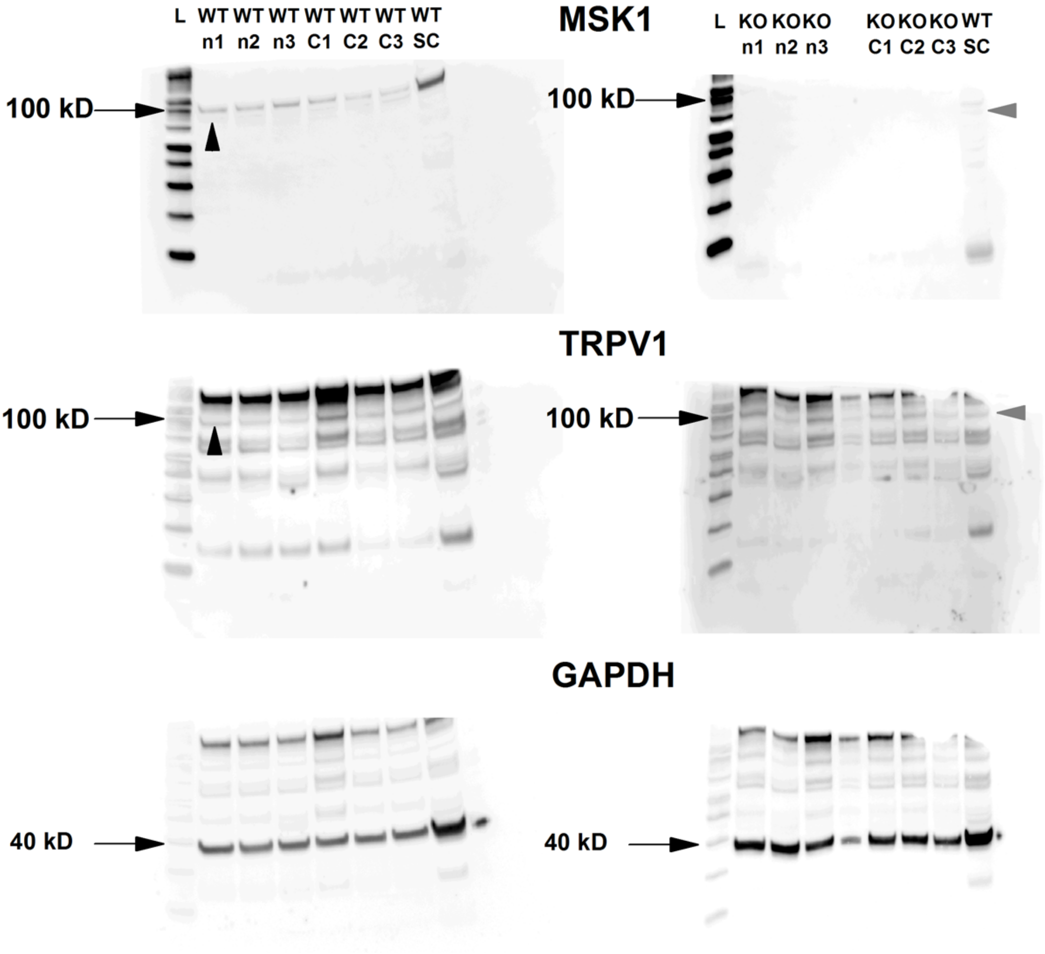


**Complete Freund's Adjuvant induces TRPV1 upregulation via MSK1**

Gel images of Western blotting of protein extracts isolated from naive (n) and CFA-injected (C) WT and MSK1$^{-/-}$ (KO) mice's L3-L5 dorsal root ganglia (DRG) 2 days after CFA injection, and a naive WT mouse's spinal cord (sc) show MSK1, TRPV1 and GAPDH expression. The lane marked by L shows weight markers. Arrowheads indicate MSK1 (MSK1 panels) and TRPV1 (TRPV1 panels) expression. Semiquantitative data are shown in Figure 6.

**Supplementary Figure 11**

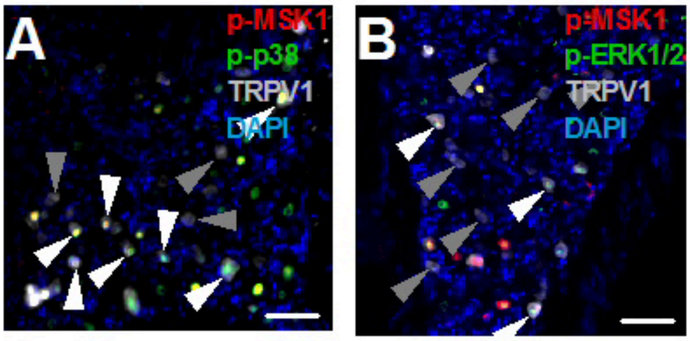


**Activated MSK1 exhibits co-expression with TRPV1 and p-p38 or p-ERK1/2**

(A) and (B) Microscopic images of WT mouse DRG sections 2 days after CFA injection into one of the hind paws showing p-MSK1, TRPV1 and p-p38 (A) or p-ERK1/2 (B) expression. White arrowheads point to TRPV1-expressing neurons which also express p-MSK1 and p-p38
(A) or p-MSK1 and p-ERK1/2 (B), whereas grey arrowheads indicate neurons expressing TRPV1 only. Scalebars indicate 50 μm.

**Supplementary Figure 12**

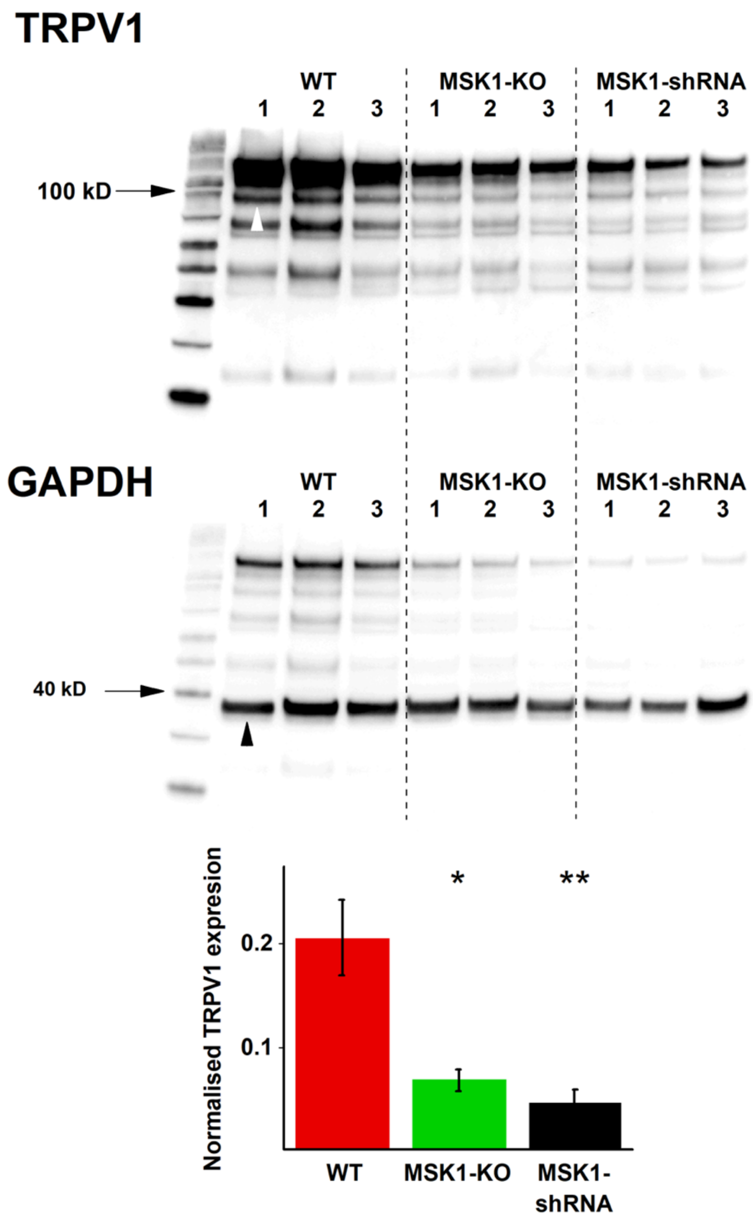


**Depletion or downregulation of MSK1 attenuates Complete Freund's Adjuvant injection-induced TRPV1 upregulation in L3-5 dorsal root ganglion**

**(A)** Gel images of Western blotting of protein extracts isolated from WT, MSK1$^{-/-}$ and AAV-*Rps6ka5*-shRNA-injected mice's L3-L5 DRG two days after CFA injection into the hind paw show TRPV1 (upper panel) and GAPDH (lower panel) expression. White and black arrowheads respectively indicate TRPV1 and GAPDH expression. **(B)** Semiquantitative analysis of gel images shown in (A) reveal significant attenuation of CFA-induced TRPV1 upregulation in MSK1$^{-/-}$ and AAV-*Rps6ka5*-shRNA-injected mice's DRG.

### Supplementary Figure 13

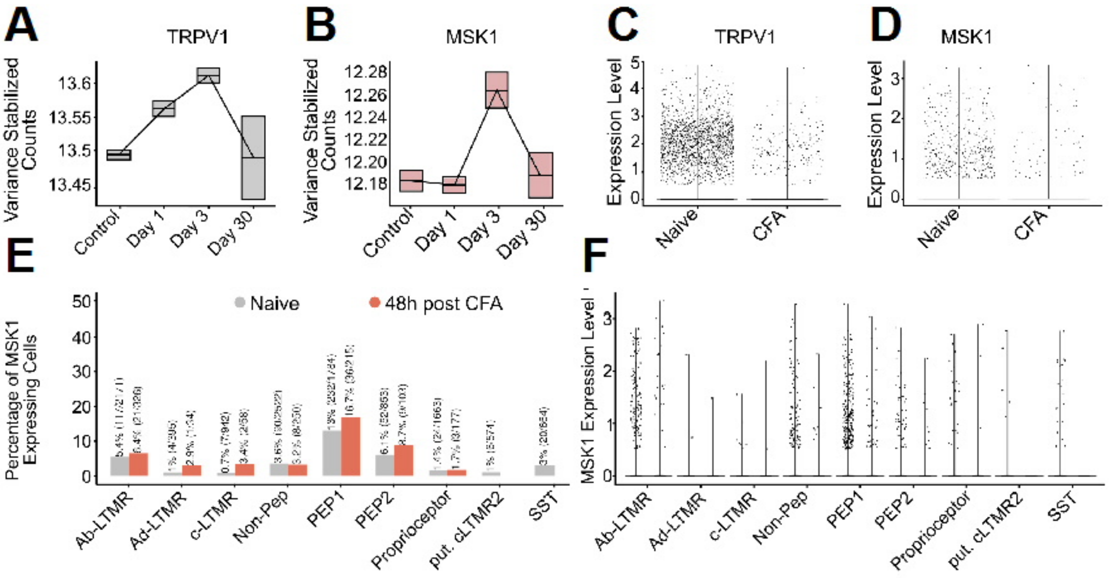


**MSK1 and TRPV1 are co-regulated by inflammation in a group of human primary sensory neurons.**

(**A**) and (**B**) Charts depict time-dependent change in *Trpv1* (TRPV1) and *Rps6ka5* (MSK1) expression in carrageenan injected mouse DRG (in silico analysis (Pierrelée et al., 2021)(GSE161764).

**(C)** and **(D)** Charts depict *Trpv1* (TRPV1; C) and *Rps6ka5* (MSK1; D) expression in mouse primary sensory neurons in naive conditions and following injecting CFA into one of the hind paws (in silico analysis (Renthal et al., 2020) GSE154659).

**(E)** Bar chart depicts the relative number of *Rps6ka5*-expresssing various transcriptionally-defined mouse primary sensory neurons in naive condition (grey) and 2 days after CFA injection into one of the hind paws (48 post-CFA, orange; in silico analysis (Renthal et al., 2020) GSE154659).

**(F)** *Rps6ka5* (MSK1) expression level in various transcriptionally-defined MSK1-expressing mouse primary sensory neurons in naive condition and 2 days after CFA injection into one of the hind paws (48 post-CFA; in silico analysis Renthal et al.: (Renthal et al., 2020) GSE154659).
