## Supplementary Table 2 for "Inflammatory burning pain depends on a specific line of nociceptors"

**Antibodies, probes and other reagents**

| **Primary antibody** | **Host** | **Supplier/ Catalogue number** | **Use** | **Dilution** |
| --- | --- | --- | --- | --- |
| Anti-MSK1 | Rabbit | Cell Signaling Technologies/3489S | Immunostaining/  Western blotting | 1:1000/ 1:1000 |
| Anti-pMSK1 | Rabbit | Cell signaling technology / 9595S | Immunostaining/  Western blotting | 1:1000/  1:1000 |
| Anti-CGRP | Goat | ABCAM/  AB36001 | Immunostaining | 1:1000 |
| Anti NF200 | Mouse | Sigma/  N5389 | Immunostaining | 1:1000 |
| Anti-NeuN | Guinea pig | Synaptic system / 266004 | Immunostaining | 1:250 |
| Anti-Trpv1 | Guinea pig | Neuromics/  403447 | Immunostaining | 1:3000 |
| Anti-Trpv1 | Mouse | Santa Cruz Biotechnology/  398417 | Western blotting | 1:500 |
| Phospho- p38  MAPK | Rabbit | Cell Signalling Technologies/  4511S | Immunostaining | 1:400 |
| Phospho- ERK1/2  MAPK | Rabbit | Cell signalling technologies/  4370S | Immunostaining | 1:600 |
| GAPDH | Mouse | Cell Signalling technologies/7076 | Western blotting | 1:1000 |
| **Secondary antibodies** | **Host/target**  **species** | **Supplier/ Catalogue number** | **Use** | **Dilution** |
| Alexa Fluor^TM^ 488  Donkey Anti Raggit IgG (H+L) | Donkey/Rabbit | Invitrogen/  A21206 | Immunostaining | 1:800 |
| Alexa Fluor^TM^ 568 Donkey Anti Rabbit IgG (H+L) | Donkey/rabbit | Invitrogen/ A10042 | Immunostaining | 1:800 |
| Cy™3 AffiniPure Fab Fragment Goat Anti-Rabbit IgG (H+L) | Goat/rabbit | Jackson Immunoresearch/  11-165-144 | Immunostaining | 1:800 |
| Alexa Fluor^TM^ 647 Goat Anti-Guinea pig IgG (H+L) | Goat/guinea pig | Invitrogen/  A21450 | Immunostaining | 1:800 |
| Alexa Fluor^TM^ 488 Donkey Anti-Guinea pig IgG (H+L) | Donkey/Guinea pig | Jackson Immunoresearch / 706-546-148 | Immunostaining | 1:800 |
| **Reagents used in histochemistry** | **Host/target**  **species** | **Supplier/ Catalogue number** | **Use** | **Dilution** |
| IB4 | n/a | Merck/  L2146 | Histochemistry | 1:1000 |
| Anti-Mouse IgG, HRP-linked antibody | n/a | Cell signaling technology / 7076(V) | Western blotting | 1:1000 |
| Anti-Rabbit IgG, HRP-linked antibody | n/a | Cell signaling technology / 7074(V) | Western blotting | 1:1000 |
| Streptavidin, Alexa Fluor^TM^ 488 conjugate | n/a | Invitrogen / S11223 | Histochemistry | 1:800 |
| **RNAscope probes** | **Host/target**  **species** | **Supplier/ Catalogue number** | **Use** | **Dilution** |
| Hs-RPS6KA5-C1 | n/a | ACDBio/1311271-C1 | In situ hybridisation | n/a |
| Mm-Trpm3 | n/a | ACDBio/Cat No. 459911 | In situ hybridisation | n/a |
| Mm-Trpa1-C3 | n/a | ACDBio/Cat No. 400211-C3 | In situ hybridisation | n/a |
| Mm-Trpv1-C2 | n/a | ACDBio/Cat No. 313331-C2 | In situ hybridisation | n/a |
